# Investigating pancreatic β cell membrane epitopes using unbiased cell-based Fab-phage display

**DOI:** 10.1101/2025.03.29.645157

**Authors:** Yena Moursli, Christian Poitras, Benoit Coulombe

**Affiliations:** Montreal Clinical Research Institute, Montréal, QC, Canada H2W 1R7; Department of biochemistry and molecular medicine, University of Montréal, QC, Canada

## Abstract

The phenotypic and functional changes of cells in response to physiological and pathological conditions are strongly influenced by the roles of plasma membrane proteins. Recombinant Fab antibody-based phage display for an unbiased antigen-driven affinity selection is a suitable approach for identifying novel membrane proteins. Alterations in the function and distribution of cell membrane proteins in pancreatic β cells have been observed in pathological conditions like diabetes. In this study, we integrated an unbiased cell-based Fab-phage display screening method with bioinformatics tools to identify and characterize Fabs that selectively bind to pancreatic β cells in conditions simulated by a hyperglycemic environment. We isolated three Fab-phages, namely Fab_53, 538, and 54.68, that have binding properties matching specific epitopes on the MIN6 membrane. These Fabs are part of the immunoglobulin G groups that contain Kappa light chains. Bioinformatics analysis of the variable domains of their light and heavy chains (VL and VH) revealed that the potential epitope binding sites on the β cell membrane are associated with pathways involved in insulin activity. Through FACS and IF analysis, we found that, of the three Fabs, Fab_538 exhibited the strongest binding to MIN6 cells. The use of InterProScan software resulted in the generation of 344 potential Fab_538 binding epitopes, and from these, AF2Complex predicted 10 interacting antigens. Our goal in combining Fab-phage display with bioinformatic tools is to develop a more effective, specific, and streamlined method for identifying disease-modifying membrane epitopes for monoclonal antibodies (mAbs).

## Introduction

The cell membrane of mammalian cells is a phospholipid bilayer that serves as an impermeable barrier to charged and hydrophobic molecules, and it contains nearly 50% proteins and 50% lipids (Cooper GM, 2000) (de Jong E., 2023). Plasma membrane proteins play key roles in ions, water, and nutrient movements across the cell membrane (Cooper GM, 2000). These proteins are also essential in shaping functional cell identity and contributing to cell heterogeneity (Stutzer I., 2012). Communication and interactions between cells in both physiological and pathological contexts also depend greatly on these cell surface proteins (Cooper GM, 2000). In fact, some studies suggest that 20 to 39% of the cell proteome consists of membrane proteins (Almén MS., 2009). However, only a limited number of these proteins have been thoroughly described and investigated, emphasizing the need to enhance our ability to screen for and identify potential new disease-modifying membrane epitopes (Aperia A., 2007).

Cell membrane proteins have become primary targets for pharmaceutical companies in biomarker discoveries and drug development due to their abnormal expression or dysfunction in various metabolic, cancer, and hereditary disorders (Cooper GM, 2000) (Köhler G., 1975). Murine mAbs were initially humanized to enable them to effectively recognize and bind to their cell membrane target epitopes while avoiding triggering an unwanted immune response. In 1975 Orthoclone OKT3 (muromonab-CD3) was one of the first monoclonal antibody drugs developed (Leavy O., 2010). This monoclonal drug was generated using hybridoma technology and was approved only 11 years later by the FDA for use in patients in 1986 (Leavy O., 2010) (Köhler G., 1975). Since then, we have seen an increase in the number of monoclonal antibodies undergoing clinical trials to reach a market value of more than 186.6 billion US dollars (USD) in 2022. This market is projected to surpass 609 billion USD by 2032, according to the Global Market Insight analysis (Faizullabhoy M., 2023).

MAbs production, as well as any form of their fragments, such as single-chain variable fragments (scFv) or fragment antigen-binding (Fab), using hybridoma technology can be time-consuming and a costly process (Alfaleh MA, 2017). On the other hand, new approaches, mainly Fab-phage display technology, are a more efficient option that allows the identification and the study of new cell membrane epitopes (Alfaleh MA, 2017). This method not only addresses the immunologic tolerance issues associated with murine antibodies but also accelerates the discovery of antigen-specific antibodies, helping to identify potential cell surface proteins that could serve as drug targets (Bradbury AR., 2011). It is a promising strategy that has successfully been used to identify new cancer cell antigens (Panagides N., 2022). Currently, 14 mAbs derived from phage display have been approved for treating various conditions, including cancers like non-small cell lung carcinoma, degenerative diseases such as macular degeneration, and autoimmune diseases like rheumatoid arthritis (Panagides N., 2022)..

Phage display leverages its extensive and diverse Fab library to enable specific affinity-based selection of antigen and epitope variants. This is made possible through the complementary amino acid interactions of the variable domains of the heavy (VH) and light (VL) chains of the Fab expressed on the phage surface with membrane proteins (Teixeira D., 2018). Using a specific Fab against a purified antigen is the most frequent method in Fab-phage display panning (Watters JM., 1997). The process involves either coating a solid surface or beads with the target molecule or using genetically modified cells that express the molecule of interest on their membrane, followed by applying the phage library to the target (McGuire MJ., 2009). However, this approach does not allow for the identification of Fabs with high cell specificity, nor does it enable the isolation of membrane proteins that distinguish between cell types or disease states (McGuire MJ., 2009). In contrast, unbiased Fab-phage library bio-panning applied to an entire cell population enables the screening and identification of antigens and epitopes in their native conformation on the cell surface when exposed to conditions that mimic a pathological state (Watters JM., 1997).

Changes in the fate and function of pancreatic β cells are key physiopathological issues in diabetes. Previous research has identified multiple β cell sub-populations, suggesting a distinct rearrangement mechanism associated with the pathological microenvironment of these cells. This, in turn, affects changes in their surface proteomic phenotype (Dorrell C., 2016). Studying β cell transmembrane proteins is challenging due to their heterogeneity, amphipathic nature, and lower abundance relative to cytoplasmic proteins (Stutzer I., 2012). Protocols that enrich and separate membrane proteins from cytosolic proteins, using subcellular fractionation or chemical labeling, followed by mass spectrometric analysis partially helped to address some of the challenges in identifying surface proteins (Elschenbroich S., 2010) (McGuire MJ., 2009). However, these methods do not capture the affinity binding sites of these proteins or the changes in their conformation in pathological contexts (Watters JM., 1997). In this study, we outline a strategy employed to identify Fabs that bind to specific cell surface proteins expressed by mouse pancreatic β cells (MIN6) using in vitro unbiased bio-panning with a Fab-phage library. The advantage of the cell-based bio-panning approach is that it enables the identification of fully functional, complex, and multimeric membrane epitopes in recapitulated pathological conditions without altering their structure through any purification process (Alfaleh MA, 2017). Thus, the primary objective of this project was to demonstrate the feasibility of isolating and identifying new Fabs and characterizing their potential specific binding sites on murine pancreatic β cells. This was achieved by combining unbiased screening of whole MIN6 cell populations with phage display and bioinformatics analysis.

The methodology presented in this study uses a subtractive approach to select cell surface antigens, receptors, or any membrane epitopes expressed by MIN6 cells in a hyperglycemic environment (figure 1). To eliminate Fab-phages binding to membrane proteins shared with other pancreatic endocrine cells and those highly abundant in various cell populations, we used murine α Tc1 clone9 cells in the initial depletion stages of the subtractive strategy. Three Fab-phage clones, Fab 53, 538 and 54.68, with distinct sequences in the variable domains of their light and heavy chains were selected in the fifth round of bio-panning. To validate the specificity of these Fabs for MIN6 cells, binding studies were conducted using Enzyme-Linked Immunosorbent Assay (ELISA), live cell sorting with flow cytometry (FACS), and immunofluorescence (IF) techniques. To bypass the time-consuming and expensive process of isolating and producing the selected three Fabs (53, 538, and 54.68) in their soluble format, we used bioinformatics software to analyze and characterize their structure and potential binding epitopes on β cell membranes.

**Figure 1:**
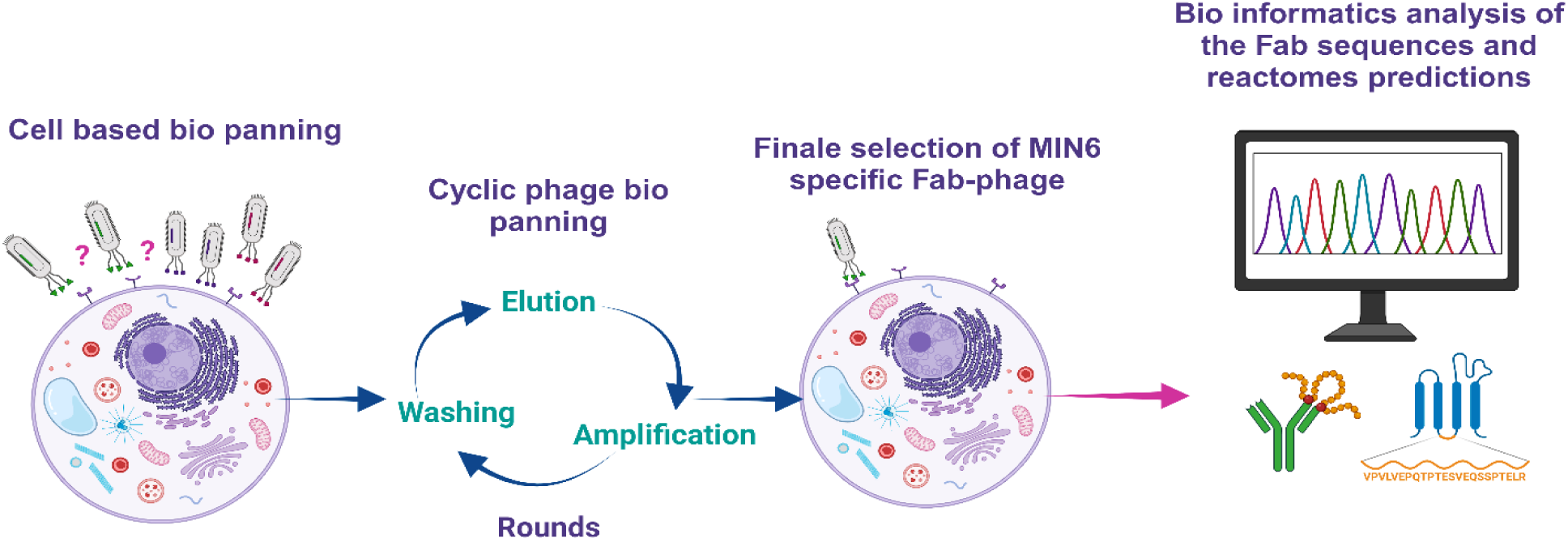
An overview of the cell-based phage biopanning experimental design. Five rounds were used to select Fabs. Bioinformatics tools, such as InterProScan and AlphaFold, were used to analyze the amino acid sequences of the Fabs and anticipate their potential binding targets on β cell surface epitopes (Figure with biorender.com).

## Materials and experiments design

### 1. Cell cultures

***The murine insulinoma pancreatic*** β ***cell, or MIN6*** cells were kindly offered by Dr. Jennifer Estall’s lab (Molecular Mechanisms of Diabetes Research Unit, Montreal Clinical Research Institute). This cell line was initially created by inducing the expression of simian virus 40T antigen in mice to obtain pancreatic tumors in transgenic mice (Poitout V, 1996). Cells were cultured in standard 100 mm petri dishes containing Dulbecco’s Modified Eagle Medium (DMEM) (1X) with 4,5 g/l of D-Glucose in 500 ml media from GIBCO (Invitrogen Corporation). The composition of the media included 15% of inactivated fetal bovine serum (FBS), 1% of antibiotic solution (5 ml of penicillin and streptomycin), L-glutamine, 110 mg/l of sodium pyruvate and 2,5 ul of 2-Mercaptoethanol (2-ME). Cells were kept in a 5 % CO2 incubator at humidified 37° C and 95 % air conditions. Every 48 to 72 hours, when cells’ confluence was at 80 to 90 %, cells were passaged. For routine cell culture maintenance, cells were detached using Trypsin-EDTA.

***The αTC1-clone 9 (aTC1-c9)*** cell line, derived from pancreatic adenomas of transgenic mice, functions as α cells that secrete glucagon. This cell line was obtained from ATCC (10801 University Blv., Manassas, VA, US, 20110-2209). These cells were created from pancreatic mice in which the SV40 large T antigen was inserted into their genome, regulated by a rat pre-proglucagon promoter. Cells were cultured in standard 100 mm petri dishes containing DMEM 1x with 3g/l of D-Glucose in 500 ml media from GIBCO (Invitrogen Corporation). The media contained 10% of inactivated FBS, 1% of antibiotic solution (5 ml of penicillin and streptomycin), L-glutamine, 110 mg/l of sodium pyruvate, 4mM of glutamine, 1% of non-essential amino acids (NEAA), 1,5% of 1M HEPES, and 1ml of 10% BSA. Cells were kept in a 10% CO2 incubator at humidified 37^0^ C and 95% air conditions. The growing medium was changed at intervals ranging from 48 to 72 hours; however, cells were passaged every 7 to 12 days depending on their confluence. For routine cell culture passages, cells were detached using Cell dissociation enzyme free Hanks’-based Buffer from Gibco.

***The NIH-3T3 fibroblast*** cell line was kindly offered to us by Dr. François Robert’s lab (Chromatin and Genomic Expression Research Unit, Montreal Clinical Research Institute). The embryonic mouse fibroblast cells were cultured in 500 ml of DMEM 1x containing 4.5g/l of D-glucose, sourced from GIBCO (Invitrogen Corporation). The media contained 10 % of FBS, 1 % of antibiotic solution (5 ml of penicillin and streptomycin), L-glutamine, 110 mg/l of sodium pyruvate, and 4mM of glutamine. Cells were kept at the same conditions as for MIN6 cells. The growing medium was changed at intervals ranging from 48 to 72 hours and passaged every 48 h. For routine cell culture passages cells were detached using Trypsin-EDTA.

Since this project focuses on identifying potential new plasma membrane epitopes by Phage-display, we chose not to use Trypsin-EDTA for cell detachment. While trypsin effectively breaks down proteins involved in cell-matrix and cell-cell interactions, it can also compromise the integrity of cell membrane proteins. Therefore, we used 0.5mM Ethylenediaminetetraacetic acid (EDTA) in sterile 1x PBS for cell detachment in all our experiments.

### 2. Unbiased cell-based bio-panning using Fab-phage display library

The unbiased cell-based bio-panning was achieved by using a premade HuFabl-5 phage display naïve human Fab library kit purchased from Creative biolabs (Creative Biolab) (Cat#: CBLX051221-2A-Unlimited Research, 45-1 Ramsy road Shirley, NY 11967 USA). The company’s original protocol was modified to meet our primary research objective of selecting an unidentified cell membrane molecule expressed at the surface of MIN6 cells.

#### 2.1. Materials and solutions

- **HuFabL-5™; the Phage Display Naïve Human Fab Library**. ∼1.0 × 10^13^ phages (∼1.0 × 10^13^ PFU/mL) supplied in 1.0 mL PBS with 10% glycerol. Complexity = 1.1 × 10^10^ transformants. Ampicillin resistant
- **E. coli TG1 Host Strain**: K12 Δ(*lac-pro*), supE, thi, hsdΔ5/F’[traD36, proAB, lacIq, lacZΔM15]. Suppressor strain, in which the amber stop codon would not be recognized
- **M13KO7 Helper Phage** at ∼1.0×10^11^ PFU phages (∼1.0 × 10^12^ PFU/mL) supplied in 0.1 mL PBS with 50% glycerol. Kanamycin resistant
- **2×TY Medium:** 1 litre of deionized H2O, 16 g/L tryptone, 10 g/L yeast extract, 5 g/L NaCl, adjust to pH 7.0 with 1 M NaOH and autoclave
- **2×TY -GA Medium**: 2×TY with 100 μg/mL ampicillin and 1 or 2 % glucose
- **2×TY -A Medium:** 2×TY with 100 μg/mL ampicillin
- **2×TY -KA Medium**: 2×TY with 100 μg/mL ampicillin, 50 μg/mL kanamycin
- **2×TY Agar**: 2×TY with 15 g/L agarose
- **2×TY Agar-AG:** 2×TY Agar with 100 μg/mL ampicillin and 1 or 2 % glucose
- **1 x Phosphate buffered saline (1 x PBS):** NaCl (137 mM), KCl (2.7 mM), Na2HPO (10 mM), KH2PO4 (1.8 mM), pH 7,4, filtered and autoclaved
- **3 % Milk 1xPBS** (3%MPBS) blocking buffer
- **Cell dissociation buffer EDTA 0.5M pH 8:** in 500 ml deionized water dissolve 93.05 grams of EDTA disodium salt (MW=372.24 g/mol) and adjust the pH with solid sodium hydroxide (NaOH) plates until pH is about 8.0, filtered and autoclaved
- **Elution Buffer:** 0.2M Glycine-HCl (pH2.2)
- **Neutralizing Buffer:** 1 M Tris, pH 7.4
- **5×PEG/NaCl:** 200 g/L polyethylene glycol-8000 (PEG-8000), 2.5 M NaCl, stir until dissolved in distilled water, filter through a 0.8 μm filter and autoclave
- **50% glycerol/1xPBS:** Equal mix of Glycerol and 1 x PBS
- **Sterile 96-well round bottom microtiter plates** made for bacterial culture (DeepWell sterile plate natural RNase/DNase-free, Thermo-scientific)
- **96-Well V-bottom microtiter plates**
- QIAprep Spin Column kit (Qiagen, Cat#: 27106)
- NanoDrop™ 2000/2000c Spectrophotometers from Thermo Scientific (Catalog number ND-2000)
- **Primers:** M13R-48: 5’-AGCGGATAACAATTTCACACAGGA-3’ (10 µM, 50 μL) and S6: 5’-GTAAATGAATTTTCTGTATGAGG-3’ (10 µM, 50 μL)
- **DNA Polymerase:** Q5® Hot Start High-Fidelity (b90275, New England Biolabs)
- **DNA Polymerase enhancer:** Q5® Hot Start High-Fidelity (b9028A, New England Biolabs)
- 5X Q5 Reaction Buffer (New England Biolabs)
- Enzyme Q5 Hot-start DNA polymerase (M0493L, New England Biolabs)
- dNTPs (10mM)
- Ladder DNA 0.5μg/ μL (10787018, Invitrogen)

#### 2.2. Key steps of a cell-based biopanning method using Fab-phage display

To identify potentially unknown epitopes on the MIN6 cell membrane, we employed a subtractive approach. These methods necessitate the use of a second, distinct cell line from the one under investigation, so we selected αTc1 for this purpose. The biopanning process typically takes 5 to 7 days per round, and each round procedure includes the following key steps:

##### 1. Depletion

In each round a depletion phase was performed by using αTc1 clone 9 cells. Cells were detached with 0.5mM EDTA solution and centrifuged for 5 min at 600 rpm. Then after counting cells, about 10 × 10^6^ cells were blocked for 1 h with 5 ml of 3% MPBS at 4°C to saturate surface antigens. In parallel, for the first round of the selection process, the phage input were prepared by aliquoting 100 µl of the phage library (containing 1 × 10^12^ phage particles) and diluting it with 900 µl of 3 % MPBS. This mixture was incubated for 1 h at 4°C on a rotator to achieve saturation. In the subsequent rounds, the phage input was adjusted based on the phage output from the previous round, aiming for 1 × 10^12^ phage particles. At the end of the 1 h, αTc1 cells were centrifuged for 5 min at 600 rpm, the blocking buffer was discarded, and the cells were resuspended in phage and 3 % MPBS mix for 2 h at 4°C.

##### 2. Selection

While incubating the αTc1 cells with the phage mix, the MIN6 cells were prepared. MIN6 cells were detached using 5 mM EDTA solution, centrifuged for 5 min at 980 rpm, and blocked for 1 h with 3% MPBS at 4°C. At the end of both MIN6 and αTc1 incubation, cells were pelleted. Then the depletion supernatant of αTc1, which contains nonbinding phages to the αTc1 membrane, was added to MIN6. MIN6 cells were incubated for 2 h with this phage mix at 4°C on a rotator.

##### 3. Washing steps

After 2 h of incubation, centrifugation was performed on MIN6 cells, and the unbound phages in the 3 % MPBS mix were discarded. Next, MIN6 cells were washed with 1 ml sterile 1x PBS. The washing steps in the first round counted for 5 times. As the number of rounds increased the number of washes increased as well. Thus, for the second round, cells were washed 10 times, and for the fourth and fifth rounds, 15 times. The unbound phages were discarded after MIN6 centrifugation for 5 min at 1100 rpm in each wash.

##### 4. Phage elution

After the final wash, the washing buffer was discarded, and 1 ml of elution buffer was added to the tube. The mixture was transferred to a protein lobind 1.5 ml Eppendorf tube and incubated for 10 min at room temperature. After 10 min, MIN6 cells were centrifuged, then the eluted phages were transferred to a new tube containing 150 µl of neutralizing buffer.

##### 5. Phage rescue

First the *E.* coli TG1 strain was cultured the same day in 2xTY medium to reach an OD600 of 0,5. However OD600 values between 0,45 and 0,55 are also accepted. After each round, 6 ml of freshly grown E. coli TG1 was added to the eluted phages in a centrifuge tube (Nalgene® Oak Ridge Style 3119 centrifuge tube) and the mix was incubated for 30 min at 37^0^ C. Next, 9 ml of 2xTY-GA medium was added to the same mix, and incubated overnight at 37°C. The next morning, the tubes were centrifuged at 5000 rpm for 15 min at 4°C. The resulting pellet was then resuspended in 1 ml of 2×TY medium and 15% glycerol and stored at −70°C.

##### 6. Phage amplification

From the rescued phage stock at −70° C, the volume of 150 µl was amplified by inoculating it into 25 ml of 2xTY-GA medium. The mixture was incubated at 37° C until it reached an OD600 value of 0.5. For the amplification, the M13KO7 helper phages was used by following the company recommendation of a ratio of 20:1 of phages to cells. Thus, 2 × 10¹¹ pfu of M13KO7 helper phages were added, and the mix was incubated at 37 degrees for 30 min. The mixture was then centrifuged during 15 min at 5000 rpm, and the pellet was resuspended in 25 ml of 2×TY-KA, followed by an overnight incubation at 30°C with shaking at 225 rpm.

##### 7. Phage purification

The following morning, the mix was first centrifuged at 4000 g for 20 min at 4^0^ C. To precipitate the phages, 5xPEG/NaCl (20%) was added to the supernatant, and the mix was incubated on ice for 1 h. At the end of the incubation, the mix was centrifuged for 20 min at 12000 g at 4^0^ C. The supernatant was discarded, and the pellet was resuspended in 1xPBS and transferred to a new 1.5 ml Eppendorf tube. To clean up the mix, a centrifugation at maximum rpm (14000) was performed. After that, the supernatant was transferred to a new, clean 1.5 ml Eppendorf tube, 20 % glycerol was added before being stored at −70°C.

##### 8. Phage titration

We conducted serial dilutions in 2xTY media to titrate eluted and amplified phage stocks from each round. We generally aimed for expected titration values of 10^12^ to 10^14^ phages/ml for amplified phages and approximately 10^8^ to 10^11^ phage/ml for eluted phages. Thus, for eluted phages the values were between 10^4^ and 10^6^, and for the amplified ones, between 10^8^ and 10^12^. Before commencing, the *E.* coli TG1 strain was grown as described previously, then 500 µL of TG1 was added to 500 µL of each phage dilution and incubated for 30 min at 37°C. The infected *E*. coli TG1 cells were spread and grown overnight at 37°C on 2xTY-GA agar plates that were previously prepared with 1 % glucose and 100 μg/ml ampicillin.

#### 2.3. Fab-Phage screening on MIN6, Fab-phages clone selection and isolation

##### Binding screening on MIN6 and αTc1

To compare binding of the isolated phages from each round to MIN6 and αTc1, the enzyme-linked immunosorbent assay technique (ELISA) was used.

##### Enzyme-Linked Immunosorbent Assay (ELISA)

To prevent cell detachment, the wells of a 96 V-bottom microtiter plate were coated with Poly-L-Lysine (Sigma Life Science, P4707). After 20 min, the Poly-L-Lysine was removed, and the wells were washed with sterile 1x PBS. Each well was then seeded with 5 to 6 × 10^4^ cells from each cell line of interest (MIN6 and αTC1). Cells were blocked with 3% MPBS for 1 h at 4°C. After 1 h, the plate was centrifuged at 900 rpm for 3 min, and the blocking buffer was removed. Concurrently, different phage-Fab titrations were prepared in 3% MPBS to be tested. Cells in the wells were then incubated with 100 µL of each phage titration for 2 h at 4°C. After 2h, cells were washed with sterile 1x PBS. To detect cell binding phages, we used an anti-M13 Major Coat Protein (RL-ph1) antibody, diluted at 1:1000 in 3% MPBS. 100 µl per well of the HRP-conjugated antibody was added and the 96 well plate was incubated at room temperature (RT) for 1h. At the end of the incubation, the wells were washed three times with 200 µl of sterile 1x PBS and incubated at RT for 5 to 20 min in darkness with 100 µl of TMB substrate. To stop the reaction, 50 µl of 10% HCl was added, and the absorbance was measured at 450 nm using a microplate reader.

##### Fab-phage clone isolation and selection

Fab-phage clone isolation was carried out following the steps outlined in figure 2.

**Figure 2:**
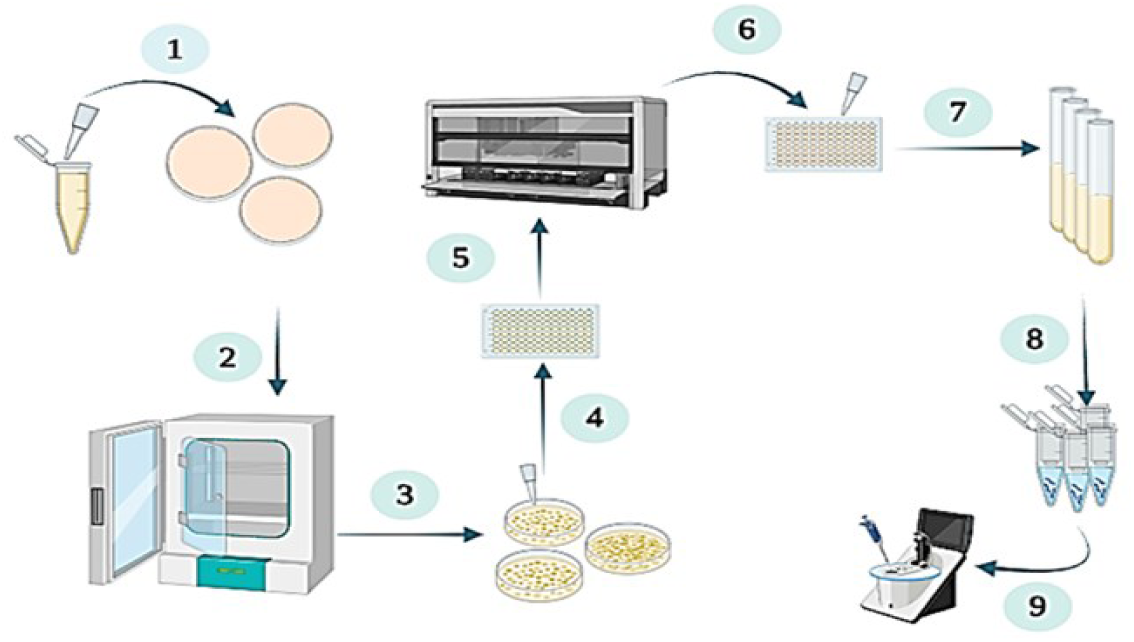
Fab-phage clone isolation and selection. 1-2) Phages from each round were smeared onto agar 2xTY-GA plates and incubated ON at 37^0^ C. 3-4) Phage plaque colonies were formed, and each colony was picked and inoculated into 100 µl of 2xTY-GA media. 5) The plate was incubated overnight at 30°C with shaking at 260 rpm. 6) A sample from each well was streaked onto a 2xTY-GA agar plate and incubated ON at 37°C. 7) A single isolated Fab-phage colony was picked and amplified in 5 ml 2xTY-A medium. 8) Phagemid-DNA extraction of each clone. 9) DNA-plasmid quantification. (Figure with biorender.com)

##### Fab-phage clone isolation

Before starting this step, agar 2xTY-GA plates were prepared in advance. Using a tip, amplified and purified phages from every round stock tube were smeared onto agar 2xTY-GA triplicate plates and incubated overnight at 37^0^ C. Each formed colony from each round titer plate was picked the next morning in order to inoculate 100 µl of 2xTY-GA media per well of a 96 DeepWell plate (DeepWell sterile plate natural RNase/DNase-free, Thermo-scientific). The plate was then incubated overnight at 30^0^ C by shaking it at 260 rpm. The next morning 40 µl of 50% glycerol was added to each well and the sealed 96 well plate was stored at −70^0^ C as a master plate for phage clones from each round of the biopanning.

##### Fab-phage DNA clone selection and isolation

From the colonies stored in the 96-well master plate, a sample from each well was streaked onto a 2xTY-GA agar plate and incubated overnight at 37^0^ C. The next morning, each single isolated Fab-phage colony from each agar plate was picked and inoculated into 5 ml of 2xTY-A medium for amplification, and the culture was incubated overnight at 37°C with shaking at 200 rpm. The following morning, before the phagemid extraction from the *E. coli* TG1, 500 µl from each overnight clone culture was removed and transferred to a 1.5 mL Eppendorf tube containing 20% glycerol and stored at −70°C for future analysis. Phagemid-DNA extraction of each clone was performed following the instructions provided with the QIAprep Spin Column kit (Qiagen, Cat#: 27106). The concentration of the final DNA-plasmid products of each Fab-phage clone from each round was measured using the NanoDrop 2000/2000c spectrophotometer device from Thermo Scientific and stored at −20°C.

##### Fab DNA amplification and verification

The Fab nucleic acid amplification was performed with polymerase chain reaction (PCR). For a total final PCR reaction volume of 50 µl, 100ng DNA plasmid was used from each sample. Forward M13R-48 and reverse S6 primers were prepared in water for a working final concentration of 10µM. The final PCR reaction master mix consisted of the following reagents from *New England Biolabs* with a final concentration of 1x for the 5X Q5 Reaction Buffer, 0.02 U/µl for Q5 Hot Start High-Fidelity DNA Polymerase and 1x for 5X Q5 High GC Enhancer. A 200 µM concentration of 10 mM dNTPs and 0.5 µM of each primer were added to the preparation.

##### PCR DNA agarose gel electrophoresis

An 0.8% (0.20 g) agarose gel was prepared using 25 ml of 1x TAE and 0.25 µg/ml ethidium bromide (EtBr). For each PCR DNA sample, 10 µl was combined with 2 µl of 5x DNA PCR buffer, and 12 µl of each sample, along with 12 µl of a DNA ladder, were loaded onto the well gel. To ensure proper separation of the PCR DNA fragments, electrophoresis was run for 45 min at a constant 100 volts.

##### Fab-DNA sequencing

After identifying positive potential Fab-DNA samples, the Fab-DNA sequences were validated using two different methods: Sanger DNA sequencing and whole plasmid sequencing. For Sanger sequencing, each Fab-DNA plasmid was prepared at a concentration of 50 ng/µl with 2 µM of the forward M13R-48 primer. For direct plasmid Fab-DNA sequencing, we followed the Plasmidsaurus sequencing protocol without the use of any primers. Thus, Fab-DNA plasmid sequencing was performed by Plasmidsaurus using Oxford Nanopore Technology with custom analysis and annotation.

#### 2.4. Bioinformatics analysis of the structures and functional properties of Fab-Phages

The identification and isolation of phage clones required the use of bioinformatics software tools to analyze and validate the Fab nature of DNA sequences.

##### Prediction design with AlphaFold software

The structure of Fab was predicted using the AlphaFold 2.3.0 version using “--model_preset=multimer” parameter. Databases required by AlphaFold were downloaded on January 17th, 2023. The structure was visualized using Mol* 3D Viewer from the Protein Data Bank Website available at https://www.rcsb.org/3d-view.

Computations were made on the supercomputer Narval, managed by Calcul Québec and the Digital Research Alliance of Canada. The operation of this supercomputer is funded by the Canada Foundation for Innovation (CFI), the Quebec ministry of economy and innovation (MEI) and the Quebec research fonds (FRQ).

This research was enabled in part by support provided by Calculation Quebec (https://www.calculquebec.ca) and the Digital Research Alliance of Canada (https://www.alliancecan.ca/).

#### 2.5. Binding assessment of the isolated Fab-phages to MIN6

To compare the binding of the identified and isolated specific Fab-phages from the most enriched rounds, we performed cell sorting using a Fluorescence-Activated Cell Sorter (FACS Aria III) and validated the results by immunofluorescence (IF).

##### Live cell sorting with flow cytometry

In their respective dishes, MIN6 and αTc1 cells were first washed with 5 ml of 1xPBS. To detach cells, 5 ml of 1xPBS and 0.5 mM EDTA were added, and then cells were incubated for 3 min at 37^0^ C. After aspiration of 1xPBS and 0.5mM EDTA, the dishes were tapped to facilitate MIN6 and aTC1 detachment. The resulting detached cells were resuspended in 5 ml of 1xPBS and centrifuged at 980 rpm for MIN6 and 550 rpm for aTC1 for 3 min. The cells were subsequently counted and aliquoted into 1.5 ml Eppendorf tubes prior to staining. For each condition, an average of 1×10^6^ to 1,5×10^6^ cells from each cell line were used for both control and experimental samples. The cells were treated and maintained at 4°C throughout the entire procedure, up to the final FACS sorting step. The identified and isolated specific Fab-phages were incubated with cells at varying PFU/ml concentrations to determine the optimal PFU/ml that produced the strongest signal. We used 3% MPBS to block the cells for 1 h, then incubated them with 200 μl of each identified Fab-phage at two different concentrations, 1×10^11^ PFU/ml and 1×10^10^ PFU/ml, for 2 h. After 2 h, the Fab-phages and cells were centrifuged and washed three times with 1xPBS for 3 minutes each. We used the anti-M13-HRP primary antibody at a 1:1000 dilution in 3% MPBS to label cells binding to phages. Cells were incubated for 1 h, then washed with 1xPBS. The Alexa 647 secondary antibody was applied at a dilution of 1:500, and the cells were incubated for 45 min before being washed three times with 1xPBS. The cells were then resuspended in 300 µl of 1xPBS, and stored at 4°C until FACS analysis, which was performed on the same day as labeling.

##### Immunofluorescence

Cells were prepared on 8-well glass slides (Lab-Tek Chamber Slide with Cover Glass Slide Sterile, Thermo Scientific, #177402) 24 h before cell staining. The well slides were coated with Poly-L-Lysine (Sigma Life Science, P4707) to guarantee cell adhesion. A volume of 200 µl was added to each well and allowed to incubate for 20 min before being aspirated. The wells were then washed three times with 1x PBS. Next, 1 × 10⁵ MIN6, αTc1, and NIH3T3 cells were seeded into each well and incubated for 24 h to allow proper adhesion to the slides before labeling. For the labeling, culture media was first removed, and cells were washed with 1xPBS then blocked with 3% MPBS for 1 h at 4°C. Phage dilutions were prepared in 3% MPBS to test two concentrations: 1 × 10¹⁰ and 1 × 10¹¹ PFU/ml. After 1 h, the blocking buffer was removed, and cells were incubated for 2 h at 4°C with phages. Cells were washed at the end of the incubation three times with 1x PBS, then fixed with 4% PFA for 15 min at RT and washed again three times with 1x PBS. We then incubated the cells overnight with the primary antibodies, anti-M13 HRP (1:1000), anti-insulin (1:500) and anti-glucagon (1:500). The next day, cells were washed three times with 1xPBS and incubated for two more hours with secondary antibodies, Alexa 647 and Alexa 488 (1:500). For NIH3T3 cell staining, Wheat Germ Agglutinin (WGA) conjugated to Alexa 488 was added at a working concentration of 5 µg/mL and cells were incubated for an additional 30 min. Finally, the slides were washed three times with 1x PBS and ProLong Gold Antifade Reagent (Thermo Fisher Scientific, #1212372) was applied. The slides were then analyzed using a Zeiss LSM700 confocal microscope.

## Results

### A subtractive approach to discover novel MIN6 surface proteins

Conventional phage display panning procedures typically involve a known target molecule, such as a purified antigen, which is either immobilized on a solid surface or genetically engineered for expression on a cell membrane. Phage libraries are subsequently directly applied to these targets. These methods are generally less complex than the challenge of identifying an unknown membrane surface epitope in its native expression. In this study, we employed a panning strategy that included a depletion step to screen for one or more unknown molecules physiologically expressed on the MIN6 cell membrane. Our methodology was inspired by two main approaches: the HuFabL-5™ Phage Display Naïve Human Fab Library Kit protocol from *Creative Biolabs* and the method developed by Yvonne Stark Y. et al. (Stark Y, 2017) as detailed in figure 3. We chose to employ a subtractive approach to identify specific proteins that were previously unknown but expressed by β cells in their physiological functional state, which involves insulin production and secretion in a high-glucose environment. The αTc1 cell line was selected for the depletion step to ensure that the binding Fab-phage targets specific MIN6 antigens, avoiding those that are highly abundant and expressed by both α and β cells.

**Figure 3:**
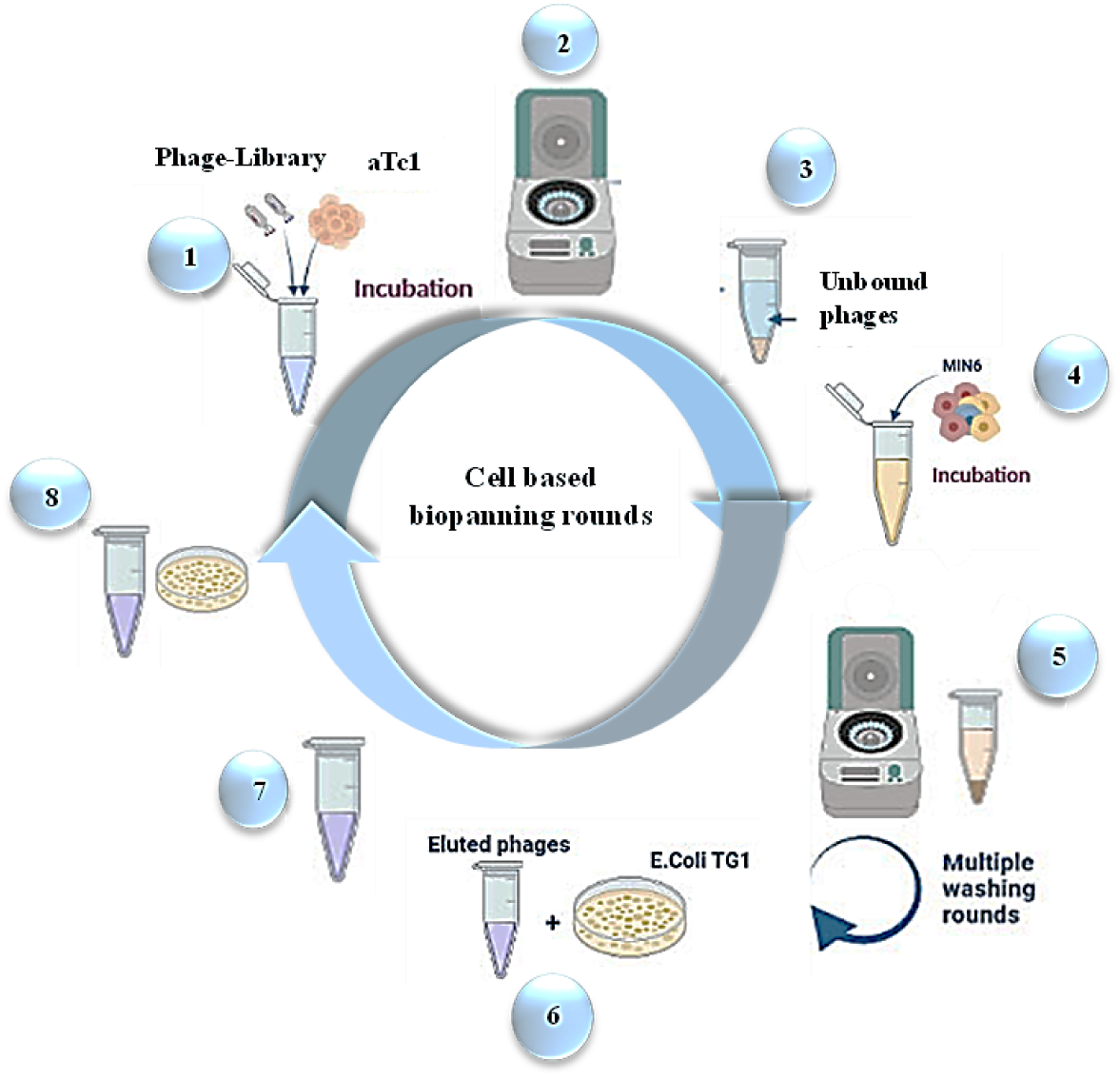
The cell-based Fab-phage display bio-panning method involves key steps in each round. 1) Subtraction: αTc1 cells are incubated for 2 h with Phages-library. 2 and 3) Unbound phages can be isolated by centrifuging αTc1 after 2 h. 4) MIN6 cells are incubated for 2 h with the unbound pool of phages. 5) Unbound phages are eliminated by washing MIN6 several times depending on the round. 6) Eluted phages are titrated to assess the enrichment level per round. 7) Eluted phages are rescued and amplified. 8) After purification phages are titrated for the next round. (Figure with biorender.com)

### The fourth and fifth biopanning rounds yielded phage clones that bound to MIN6

In total, we performed six rounds of depletion, selection, phage elution/rescue/amplification/purification, and titration. We then screened for specific Fab-phage binding to MIN6 through multiple rounds using an ELISA. The fifth round yielded the highest output titers, indicating late enrichment and suggesting that the positive, specific phages were more abundant in the fifth round than in the third or fourth rounds. The enrichment factor between rounds increased after the third round of binding selection, rising from 0.6 in the third round to 2 in the fourth round and a peak of 103.7 in the fifth round (figure 4a). To determine if the phage pools from the fifth and fourth rounds of cell-based biopanning exhibited a stronger specific binding for MIN6 than for αTc1, we performed a polyclonal ELISA (figure 4b). While no significant increase in enrichment was observed in the second and third rounds, the phage pools showed a stronger binding to αTc1 than to MIN6. However, in the fourth and fifth rounds, where enrichment levels significantly increased, the binding of the phages to MIN6 was 4 times higher than for αTc1 in the fourth round and 2.4 times higher in the fifth round (figure 4b).

**Figure 4:**
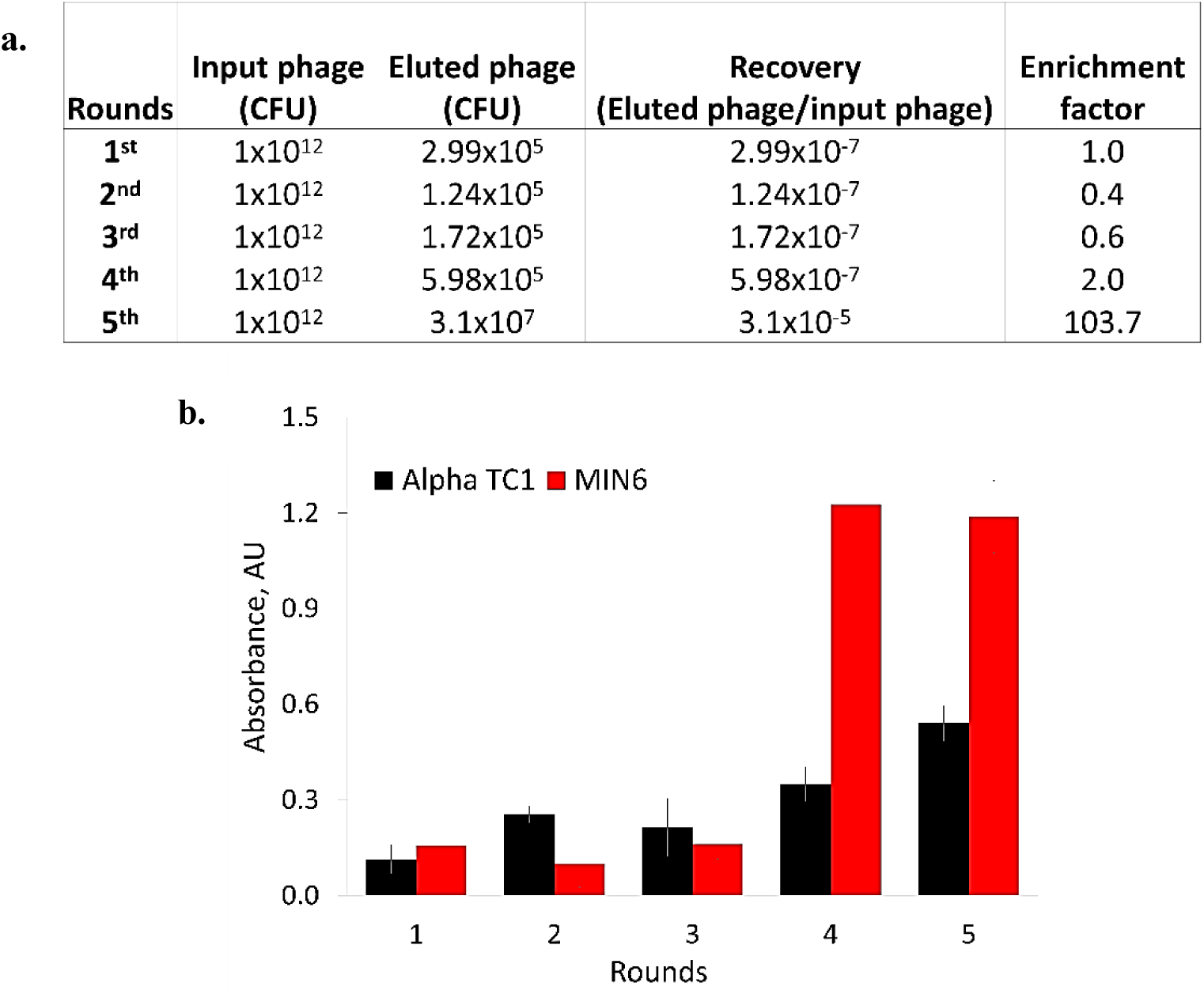
Assessment results of phage binding to MIN6 by ELISA between rounds. a) Five rounds of bio-panning resulted in the enrichment of MIN6 cell specific phages. The fourth and fifth rounds of bio-panning led to a marked increase in the number of phage particles specific to MIN6 cells. b) ELISA results obtained using the amplified and purified phage eluates from each round of bio-panning. Phages binding to MIN6 and αTc1 cells were assessed using HRP-conjugated anti-M13 phage antibody. ELISA revealed that MIN6 cells had higher absorbance values compared to αTc1cells during rounds four and five, suggesting that the phage pools have a higher affinity to MIN6 cells.

### Identification of three potential Fab-phages with sequence variability in their variable domains

After isolating, amplifying, and purifying the Fab-phage clones from the third, fourth, and fifth rounds, we employed two methods—Sanger sequencing and whole plasmid sequencing —to validate the Fab-DNA sequences. PCR-DNA agarose gel band results were used to select 59 Fab-phage DNA clones, primarily from the fifth round, for sequencing.

We decided to combine two sequencing methods for the following reasons. First, to ensure accuracy, as bacterial hosts like *E. coli* often undergo significant gene mutations to avoid expressing genes that may be toxic to the host. These genetic modifications in the bacterial genome could also affect the plasmid being studied in unforeseen ways (Rosano GL., 2014.). When using a targeted sequencing method like Sanger sequencing, these altered sequences may go undetected. Therefore, it is essential to sequence the entire plasmid to confirm that all sections of the Fab-phage are present before concluding that the obtained sequences correspond to a potential Fab-phage. The second reason is cost-efficiency. Due to the length and complexity of the plasmid sequences in our samples, it was more economical to sequence the entire plasmid upfront rather than performing multiple Sanger sequencing runs on the same plasmid (figure 5).

**Figure 5:**
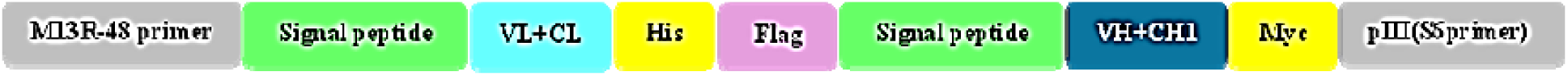
Diagram of the composition of the Fabs plasmid DNA sections. The diagram shows the structure of the Fab-phage library (HuFabL-5™ from Creative Biolab) used in cell-based biopannig. It highlights the expression cassette of psd-hu FabL-5, including the plasmid sections that were sequenced and amplified using PCR, Sanger sequencing, and Plasmidsaurus. These sections include the forward primer M13R-48 region, signal peptide, variant and constant light chain sequences of the Fab, His tag, Flag tag, variable and constant heavy chains, Myc tag, and the reverse S6 protein primer.

Upon analyzing the whole Fab-DNA plasmid sequencing results, we confirmed that the plasmid structure cassette provided by the manufacturer was correctly incorporated into our samples (figure 5). Most of the analyzed sequences showed similarity to Fab_53, which was consistent with the results from several agarose DNA-PCR gels. Only Fabs_538 and 54.68 displayed differences from Fab_53 in both the whole Fab-DNA plasmid (figure 6) and Sanger sequencing results (figure 7a).

**Figure 6:**
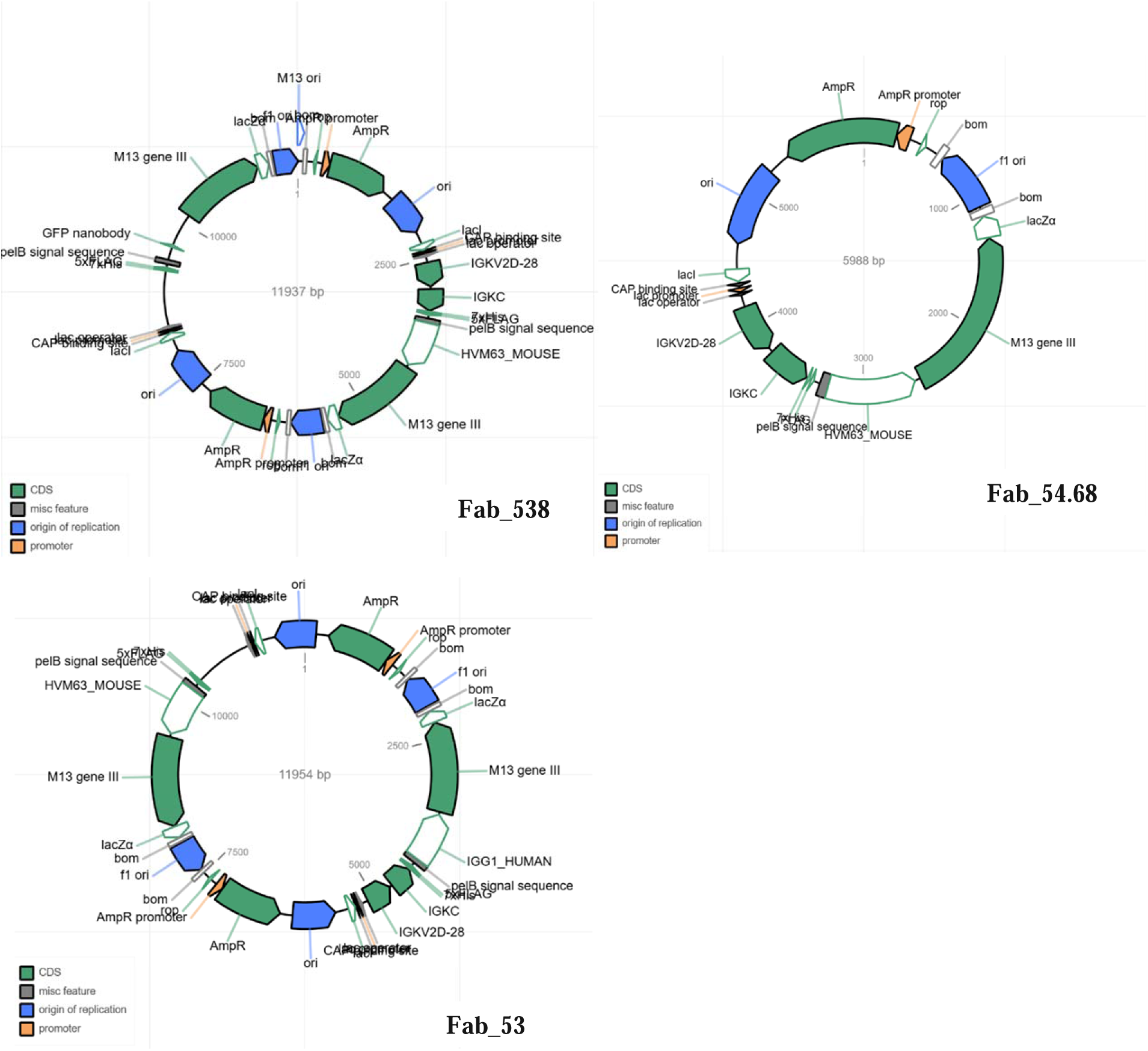
Plasmidsaurus sequence analysis of the three Fab plasmid DNAs. The sequencing and analysis results of the Fabs_53, 538, and 54.68 plasmids are presented, with each section clearly marked: the coding sequence (CDS) is green, the miscellaneous features in grey, the origin of replication in blue, and the promoter region in orange. These three plasmid images were generated by Plasmidsaurus.

**Figure 7:**
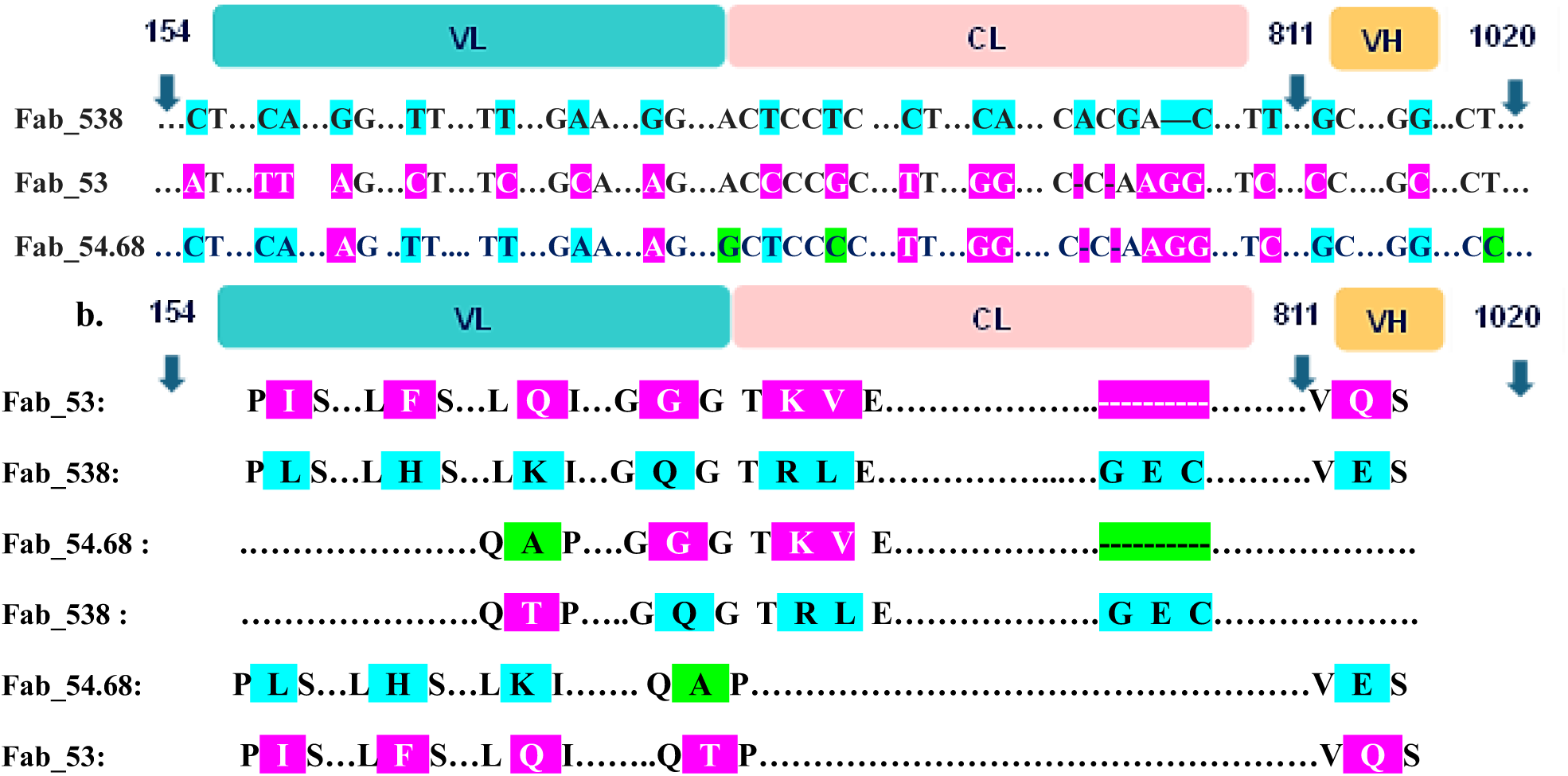
Comparison of nucleotide and amino acid sequences of the VL, CL, and VH domains between the three Fabs. a) A comparison of the DNA sequences of the light and heavy chains of Fabs 53, 538, and 54.68 to determine their differences in the variable and constant regions. Blue and purple are used to distinguish Fab_538 from Fab_53, and green highlights differences between Fab_54.68 and the other two Fabs. b) A comparison of the translated protein sequences of Fabs 53, 538, and 54.68 highlighting differences mostly in the VL domains.

The Fab_538 light chain, comprising both variable and constant regions, has the longest protein length at 219 amino acids, compared to 216 for Fab_53 and Fab_54.68. The VL+CL protein sequences of Fab_538 share 214 amino acids with those of Fab_53 and Fab_54.68, exhibiting 97.72 % similarity. The VL+CL sequences of Fab_53 and Fab_54.68 show a 99.07 % similarity, as 214 out of 216 amino acids are identical. Variations in the amino acid sequences between Fabs_538 and 53 are mainly found in the variable and constant domains of their light chains, particularly in the first 110 amino acids. Differences were also observed between the Fabs_53 and 54,68 within the first 100 amino acids (figure 7a and b). Interestingly, although the VL+CL regions of Fab_538 showed equivalent similarity to both Fab_53 and Fab_54.68, there were fewer differences between Fab_538 and Fab_54.68 compared to Fab_538 and Fab_53 (figure 7b.)

Using sequence nucleotide alignment software, we analyzed the amino acid variations among the three Fabs by comparing their open reading frames (ORFs). We examined three ORFs, and the notable nucleotide variations resulted in amino acid differences between the three Fabs, particularly in the longest first frame (figure 1 in supplementary material). These variations were primarily located within the 0 to 360 nucleotide regions of the VL, the last 9 nucleotides of the CL, and the first 25 nucleotides of the VH (figure 7b.). Fab_53 is distinct from Fab_538 by six specific amino acids, as detailed in Table 1.

**Table 1:** Table of amino acid variations in the VL and CL domains across Fabs. The following table presents the amino acid variations between Fabs in the VL and CL domains. Amino acids shared between Fab_53 and the other two Fabs (Fab_538 and Fab_54.68) are highlighted in purple to represent Fab_53. Amino acids in Fab_538 that differ from those of Fab_53 at the same position are highlighted in blue, and these differences are also shared with Fab_54.68. The amino acid (alanine) at position 96 in Fab_54.68 is highlighted in green, as it is not shared with either Fab_53 or Fab_538.

| Fab chains | Variation in residues between the Fabs |  |  | Locations |
| --- | --- | --- | --- | --- |
| VL | **Fab_53** | **Fab_538** | **Fab_54.68** |  |
|  | Isoleucine (I) | Leucine (L) | Leucine (L) | 6 |
|  | Phenylalanine (F) | Histidine (H) | Histidine (H) | 28 |
|  | Glutamine (Q) | Lysine (K) | Lysine (K) | 76 |
|  | Threonine (T) | Threonine (T) | Alanine (A) | 96 |
|  | Glycine (G) | Glutamine (Q) | Glycine (G) | 102 |
|  | Lysine (K) | Arginine (R) | Lysine (K) | 105 |
|  | Valine (V) | Leucine (L) | Valine (V) | 106 |
| CL |  | Glycine (G) |  | 214 |
|  |  | Glutamic acid (E) |  | 215 |
|  |  | Cysteine (C) |  | 216 |

### Validation of the structure of the three identified Fabs using AlphaFold computational analysis

Since the Fabs_53, 538 and 54.68 have not been previously described or identified in any prior studies or experiments, we used AlphaFold 2 to confirm that these molecules were indeed Fabs and not other types of protein groups. Using the primary amino acid sequences of the variable and constant domains of the light and heavy chains of the Fabs, AlphaFold was able to predict their 3D structures (figure 8). The amino acids defining the specific features of the VL+CL domain of all three Fabs_53, 538 and 54.68 in the AlphaFold-predicted structure align with our comparative analysis presented in Table 1. For Fab_53, the defining amino acids include Isoleucine (I), Phenylalanine (F), Glutamine (Q), Threonine (T), Glycine (G), Lysine (K), and Valine (V). For Fab_538, they are Leucine (L), Histidine (H), Lysine (K), Threonine (T), Glutamine (Q), Arginine (R), Leucine (L), Glycine (G), Glutamic acid (E), and Cysteine (C). For Fab_54.68, the amino acids include Leucine (L), Histidine (H), Lysine (K), Alanine (A), Glycine (G), Lysine (K), and Valine (V).

**Figure 8:**
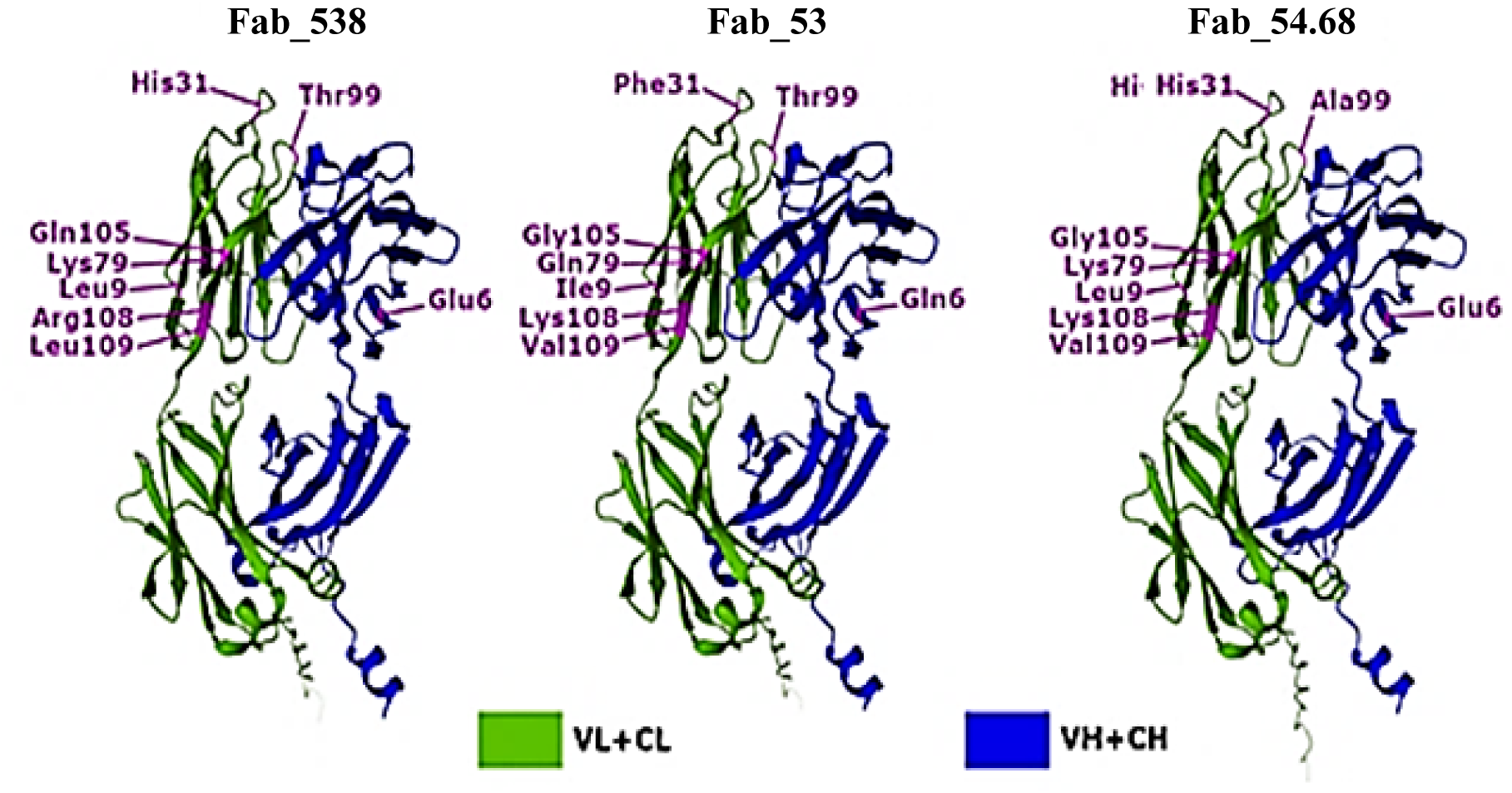
2D figures of the 3D structures of the three Fabs generated with AlphaFold. 2D figures of the AlphaFold predicted 3D structure of the Fabs_53, 538 and 54.68. The variable and constant light chain domains (VL+CL) are shown in green, while the heavy chain domains (VH+CH) are depicted in blue. The purple color highlights the location of amino acids within the VL+CL and VH+CH domains.

We also found that, unlike the VL+CL domains of the Fabs, which exhibit several amino acids that characterize them, the VH+CH domain primarily displayed a single variable amino acid among the three Fabs: Glutamine (Q) for Fab_53 and Glutamic acid (E) for both Fab_538 and Fab_54.68 (figure 8).

### Fab_538 showed stronger binding to MIN6

FACS analysis was performed to evaluate whether the three Fab-phages, Fab_53, 538, and 54.68, isolated from the fifth round of biopanning, exhibited strong binding to MIN6 compared to αTc1 cells. MIN6 and αTc1 cells were incubated for 2 h with two different concentrations of each Fab-phage: 1 × 10¹⁰ and 1 × 10¹¹ PFU/ml. Control groups included MIN6 and αTc1 cells incubated with only the primary and secondary antibodies, without any phages. Compared to αTc1 and the control groups, a significantly higher number of MIN6 cells bound to Fab_538 at both concentrations (figure 9a). Although Fab_53 and 54.68 demonstrated a stronger binding to MIN6 compared to αTc1, it was still lower than the specific binding observed with Fab_538. In three independent analyses, the average percentage of MIN6 cells expressing a membrane protein specifically binding to Fab_538 was 28.8 %, compared to 8.9% for αTc1 (figure 9b).

**Figure 9:**
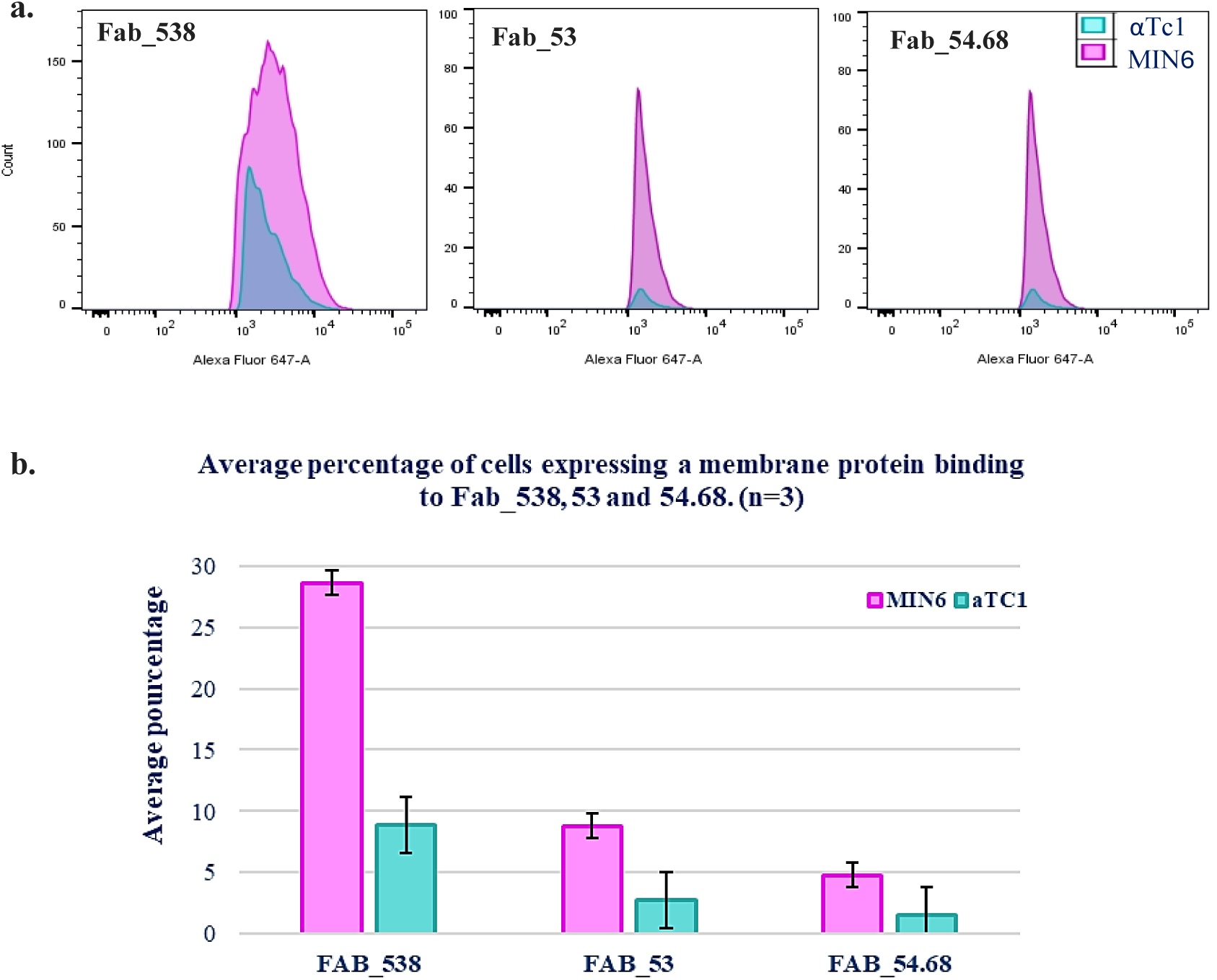
FACS binding analysis of the three Fabs to MIN6 cells. Comparative FACS analysis of the specific binding of the three identified Fabs-phages (53, 538, and 54.68) to MIN6 and αTc1 cells. Fab-phage binding was assessed using a primary mouse anti-M13 antibody, followed by a secondary anti-mouse Alexa 647 antibody. a) At a Fab-phage concentration of 1 × 10¹¹ PFU/ml, a distinct signal shift is observed, with a higher number of MIN6-M13+ cells specifically binding to Fab_538 compared to αTc1 cells. Although Fab_53 and Fab_54.68 also exhibit specific binding to MIN6 cells over αTc1, their binding intensity is lower than that of Fab_538. b) 28.8% of MIN6 cells are Fab_538-M13+ compared to 8.9% of αTc1 cells, the affinity binding to MIN6 is also higher with Fab_538 than with Fabs_53 and 54.68.

### Immunofluorescence assay confirming the strong binding of Fab_538 to MIN6 cells

To validate the strong specific binding of Fab_538 to MIN6, we compared it with a cell line different from pancreatic endocrine cells. The NIH-3T3 cell line was incubated with Fab_538-phage for 2 h, alongside MIN6 and αTc1 cells. Primary antibodies targeting insulin and glucagon, the two well-known hormonal markers of b and a cells, were used to differentiate the two cell types. Wheat Germ Agglutinin (WGA) conjugated to Alexa Fluor 488 labeled the plasma membrane of the NIH-3T3 mouse fibroblast cells (figure 10 a and b).

**Figure 10:**
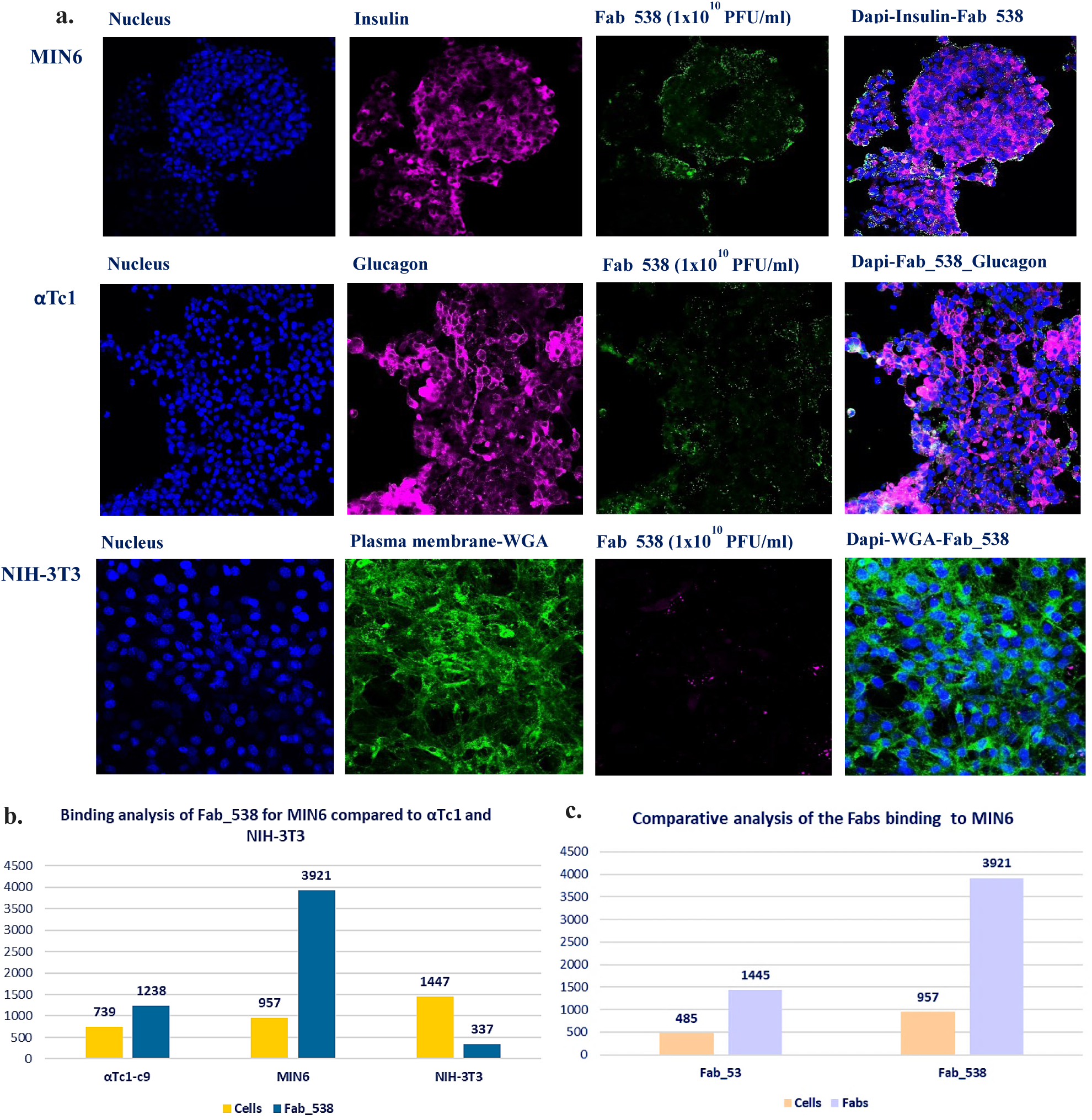
Immunofluorescence validation of Fab_538 binding to MIN6. a) The binding specificity of Fab_538 to MIN6 cells was confirmed through IF. Blue represents DAPI staining for the nucleus, purple indicates Alexa 647 staining for insulin in MIN6, glucagon in αTc1, and Fab_538 in NIH-3T3 cells. Green represents Alexa 488 staining for Fab_538 in MIN6 and αTc1 cells, as well as WGA labeling of the plasma membrane in NIH-3T3 cells. b) ImageJ intensity analysis software was used to estimate the binding of Phage-Fab_538 to individual cell types and the cell count per image. The strong specific binding of Fab_538 to MIN6 was confirmed, with 3921 phages binding to 957 cells, compared to 1238 phages binding to 739 αTc1 cells. This indicates a binding rate 4 times higher for MIN6 than for αTc1, with a ratio of 1.6 for αTc1. Additionally, the affinity of Fab_538 for MIN6 was further validated by comparing it to the phage’s binding for NIH-3T3, which showed a much lower ratio of 0.23. c) ImageJ intensity-based comparative analysis of the immunofluorescence revealed that Fab_53 has a lower binding for MIN6 compared to Fab_538, with almost half as many phages binding to MIN6.

The binding of Fab_538 to MIN6, compared to αTc1 and NIH-3T3 cells, was confirmed by observing 3921 phages binding to 957 MIN6 cells, corresponding to a ratio of 4 phages per cell. In contrast, only 1238 phages bound to 739 αTc1cells, resulting in a ratio of 1.6 phages per αTc1 cell. We also assessed whether Fab_538 binds to a cell line unrelated to pancreatic Langerhans islet cells. The NIH-3T3 (mouse fibroblast) binding assay for Fab_538 showed that only 337 phages bound to 1447 cells, resulting in less than one phage per cell (a ratio of 0.23) (figure 10a. and 10b.). Comparative binding analysis showed that Fab_538 binds more strongly to MIN6 than Fab_53, with 4 phages of Fab_538 binding to MIN6 compared to 2.9 phages of Fab_53 (figure 10c).

### The binding of Fab_538 to MIN6 cells increases in medium containing 1 g/L glucose

All rounds of Fab-phage display biopanning were performed with MIN6 cells cultured in medium containing 4.5 g/l glucose. Therefore, the three isolated Fabs, namely Fab_538, Fab_53, and Fab_53.68, are capable of binding to membrane proteins expressed by MIN6 cells in a high-glucose environment. Additionally, InterProScan analysis of the potential reactomes suggested that these Fabs may interact with receptors associated with the insulin signaling pathway. To assess the functional variation in the binding of these Fabs to MIN6 cells based on glucose concentration, MIN6 cells were cultured for 48 h in DMEM with 1 g/l glucose and incubated with Fabs_53, Fab_538, and Fab_54.68. ELISA results comparing the binding of these Fabs to MIN6 at different glucose concentrations showed that the binding of all Fabs increased after 48 h of culture in 1 g/l glucose (figure 11). Notably, binding of Fab_538 to MIN6 increased by 90 % (figure 11).

**Figure 11:**
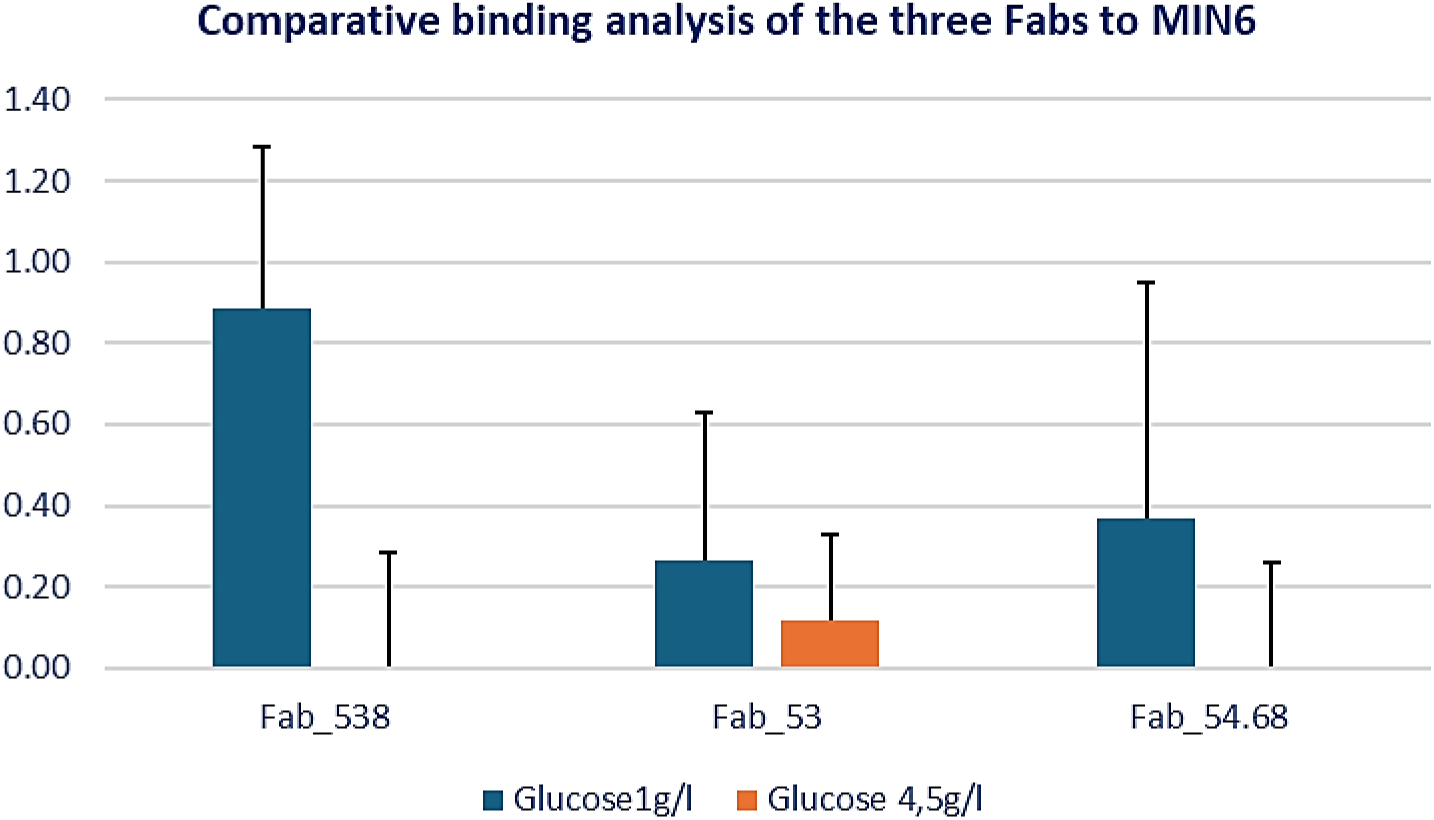
ELISA analysis of Fabs binding to MIN6 cells based on glucose concentration in the medium. ELISA signal results at 450 nm of the isolated Fab-phages (Fab_538, 53, and 54.68) at 1×10^10^ PFU/ml, binding to MIN6 cells cultured for 48 h in DMEM with 1 g/l and 4.5 g/l glucose concentrations. In a 96-well plate, 10^5^ MIN6 cells were seeded per well (three wells per condition and per Fab), blocked with 3% MPBS for 1 h, and then incubated for 2 h with each Fab. After washing off unbound phages, an anti-M13 coat protein HRP antibody was used to detect phage binding to MIN6. Control wells consisted of MIN6 cells cultured in DMEM with 1 g/l and 4.5 g/l glucose concentrations, but without phage incubation.

### InterProScan analysis confirmed that the three Fabs belong to the IgG class

A major challenge in unbiased screening for new, potentially functional cell membrane epitopes using a Fab-phage library is the expensive and time-consuming process of isolating and purifying soluble Fabs from selected phages, followed by conducting experimental functional assays and characterization. These challenges are further exacerbated when the isolated Fabs-phages may not actually be Fabs. Therefore, it is essential to use advanced computational tools and databases to confirm that the identified Fabs belong to an existing immunoglobulin family. To address this challenge, InterProScan was used as a comprehensive bioinformatics tool, to analyze the longest open reading frame amino acid sequences of VL, CL, VH and CH from the three Fabs—538, 53, and 54.68.

InterProScan analysis results confirmed that both VL and CL sequences are part of the immunoglobulin (Ig) light chain families. Specifically, the variable and constant domains of all three Fabs belong to the kappa light chain. Immunoglobulin are classified into five major types— IgG, IgM, IgA, IgE, and IgD—based on the respective constant domain of their heavy chains: γ, μ, α, ε, and δ (Schroeder HW Jr, 2010). The constant domains of the heavy chains in the three Fabs aligned with the IgG sequences, according to the results from FUNFAM database (figures 3, 4 and 5 in supplementary materials).

### Prediction of the functional epitopes of the Fab_538

Among the three Fabs, Fab_538 demonstrated the strongest binding to MIN6 cells under both high and normal glucose concentrations in vitro. As a result, we focused our analysis on the antigen-binding site of Fab_538 to pinpoint potential functional binding sites. To explore possible reactomes, we utilized InterProScan to compare the amino acid sequences of the VL domain with other known immunoglobulin sequences. InterProScan uses profile scan-match across multiple databases, integrating data from sources like SUPERFAMILY, PANTHER, PROSITE, CATH-Gene3D, PIRSF, the Conserved Domains Database (CDD), PRINTS, HAMAP, NCBI, Pfam, SMART, and the Structure-Function Linkage Database (SFLD) (Blum M, 2025).

The antigen-binding sites of immunoglobulins (Igs) are traditionally defined by the combination of the three complementarity-determining regions (CDRs) from the variable light (VL) and variable heavy (VH) domains (Schroeder HW Jr, 2010). InterProScan data identified six Ig domain chains corresponding to the VL sequences of Fab_538 (Table 1 in the supplementary materials). These six Igs include the Ig-like domain superfamily, the Ig V-set domain, the Ig-like fold, the Ig-domain subtype, and both the Ig-heavy variable and Ig-variable light chain domains. The analysis of the extensive list of potential cell surface-interacting antigens generated by InterProScan was focused on the reactome identifiers of *Mus musculus*. A total of 344 potential Fab_538 binding epitopes were selected, including receptors associated with the insulin pathway. These processes include insulin receptor recycling, the insulin receptor signaling cascade, insulin processing, and regulating Insulin-like Growth Factor (IGF) transport and uptake by Insulin-like Growth Factor Binding Proteins (IGFBPs).

To predict the potential binding sites of Fab_538, AF2Complex uses computational models to analyze specific amino acid residues within the antibody’s complementarity-determining regions (CDRs) most likely involved in binding. These predictions are based on the 344 epitopes generated using InterProscan, which help identify residues in the antibody’s CDR loops. AF2Complex’s structure prediction accurately identifies the precise positioning of amino acids in the variable domains of Fab_538. AF2Complex algorithms accurately predict the 3D structure of these regions and identify the amino acids in the CDRs with high probability to interact with the antigen and translate the results into an interaction score of potential binding sites of the antibody (Polonsky K., 2023). Thus, out of the 344 epitopes, AF2Complex’s predicted 650 protein targets for complementary binding to the VL of Fab_538. From these 650 proteins, we retained 10 proteins with an interface score greater than 0.5 (figure 12 a.), suggesting a potential for Fab_538 to bind to MIN6 cells. The number 1 represents the perfect interface score (IS) suggesting that the predicted protein has a 100 % chance of structurally docking into the CDR binding region of the Fab_538, and 0 has the least chance of being a binding site. Thus, we decided, based on Abramson J. et *al.* outcomes to select the proteins with a score greater than 0.5 which indicates that the ligands have a 50 % or more probability of binding to Fab_538 (Gao M, 2022). Among these proteins, three have been described in β cells, while seven have not yet been identified or characterized in β cells. Additionally, five are known to be expressed on the plasma membrane (figure 12 b.)

**Figure 12:**
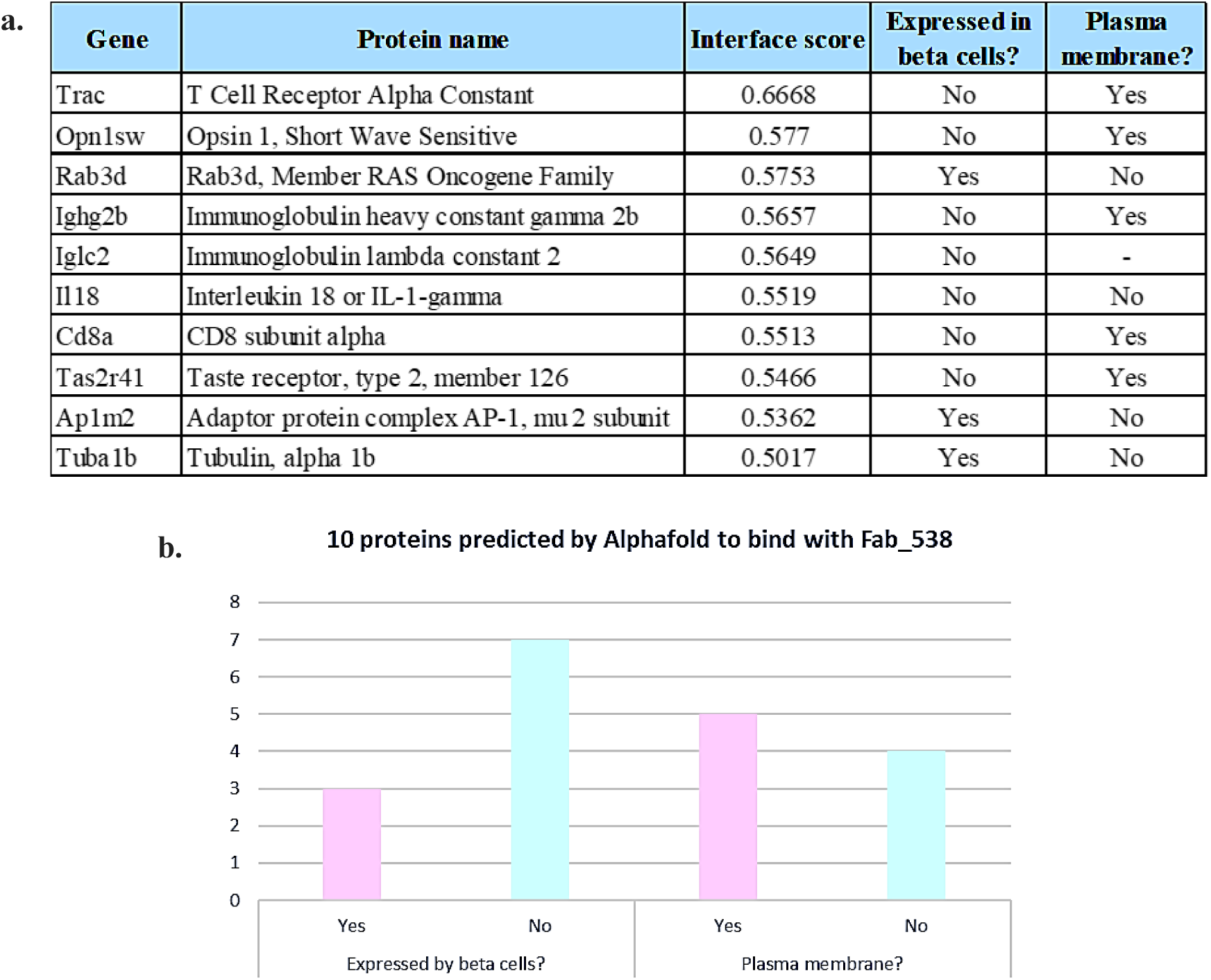
List of the 10 predicted ligand proteins for Fab_538 identified by AF2Complex. a.) 10 proteins with the potential to bind to the variable domains of Fab_538, each having an interface score greater than 0.5, were selected. b.) Of the 10 proteins, 3 are described as being expressed by pancreatic β cells, while 7 have not been identified in β cells. Additionally, only 5 of the 10 proteins are known to be expressed on the plasma membrane.

## Discussion and limitations

Diabetes mellitus (DM) is a chronic metabolic disorder usually associated with several other life-threatening conditions such as stroke, myocardial infarction or renal insufficiency that are all the consequence of chronic glucose homeostasis failure (Kottaisamy CPD., 2021). Currently, two types of DM have been described based on the underlying pathophysiological mechanisms, clinical progression, and the affected patient groups. Type 1 DM (T1DM) which is an autoimmune disorder commonly affecting children and young adults (Marro BS., 2017) and type 2 DM (T2DM), a non-insulin dependent form of diabetes (Rubio-Navarro A., 2023). In both type 1 and 2 diabetes, and regardless of the underlying pathophysiological processes, insulin production by β cells is always compromised.

1975 marked the discovery of the first monoclonal antibody (mAb) by Kohler and Milstein, which has led to a new era of using mAbs in the treatment of various diseases (Köhler G., 1975). However, the entire conventional process—from identifying the appropriate cell membrane protein or therapeutic target to the production and testing of humanized mAbs—is exceptionally long, complex, and expensive (Berger M., 2002). One of the most traditional approaches used to generate mAbs is the hybridoma technology, which involves fusing spleen cells from humanized mice with murine myeloma cells (Ke Q., 2021). This method was used to develop the first humanized mAb, Teplizumab, which targets CD3 on T lymphocytes, for type 1 diabetes treatment (Masharani UB, 2010). It was developed in 1995 but was approved until November 2022 by the Food and Drug Administration (FDA) for patients with type 1 diabetes aged over 8 years (Ramos EL., 2023) (Keam SJ., 2023),.

In contrast, phage display technology has made the screening and identification of antibodies targeting cell membrane epitopes much more convenient. Phage display, particularly using filamentous bacteriophage M13, has gained popularity for producing recombinant antibody fragments (Fabs) to search for new cell surface antigens, epitopes, or receptors (Bazan J, 2012) (Ljungars A, 2019). The M13 bacteriophage genome consists of a circular, single-stranded DNA molecule enclosed by a coat protein called pVIII, which makes up 98% of all M13 coat proteins, along with additional coat proteins, including the pIII and the pVII + pIX complex (Saw PE, 2019). Usually, the Fab displayed by the M13 phage is fused to the pIII (Saw PE, 2019).

Phage display panning technology is commonly used by exposing phages to soluble or immobilized recombinant proteins or antigens (Bradbury AR., 2011). The purity and quality of the antigens used are essential for the success of this approach (Jones ML, 2016.). But these methods have the drawback of not being able to identify novel or unknown cell surface antigens or epitopes in their native conformations (Alfaleh MA, 2017) (Stark Y, 2017). Because transmembrane proteins are tightly bound to the phospholipid bilayer, it has been demonstrated that using dispersive agents, such as detergents, to extract them alter their structure and physiological functions (Tao X, 2023.). On the other hand, unbiased whole cell biopanning procedures are suitable for the identification of cell membrane proteins in their native physiological structure (Alfaleh MA, 2017) (Stark Y, 2017).

Cell-based Fab-phage display biopanning presents some challenges, such as removing non-specific cell surface binders and isolating and purifying soluble Fab from the phage. Nevertheless, it remains one of the most effective methods for identifying cell surface epitopes or antigens in their natural configuration, including after undergoing post-translational modifications (Sánchez-Martín D, 2015). In this proof-of-concept project, we used unbiased whole pancreatic murine β cells (MIN6) as the target for *in vitro* Fab-phage display bio-panning, with αTc1-Clone 9 serving as the subtractive cell line.

To study pancreatic β cells *in vitro* and understand how their functions change in a diabetic context, media with glucose concentrations that mimic hyperglycemic conditions are often used. Typically, a normoglycemic glucose concentration of 5.5 mM is used *in vitro,* while glucose concentrations ranging from 25 to 50 mM (4,5g/l to 9g/l) are commonly used to replicate diabetic conditions (Aldoss A., 2023). In our study, for the cell-based phage biopanning, MIN6 and αTc1 cells were cultured in media with high glucose concentrations: 4.5 g/l for MIN6 and 3 g/l for αTc1.

To identify a pool of phages specifically binding to MIN6 and αTc1 cells during the 3rd, 4th, 5th, and 6th rounds of biopanning, we employed the ELISA method. This technique is fast and cost-effective for screening Fab-phage clones with specific binding to MIN6. The results revealed that MIN6 cells exhibited higher absorbance values than αTc1 cells in rounds four and five, indicating a stronger binding of the phage pools to MIN6 cells (figure 4). Based on these results, 48 Fab-phage clones from the third round, 45 from the fourth round, and 48 from the fifth round were isolated. Despite isolating a significant number of Fab-phage clones, DNA analysis and sequencing revealed only three Fabs—53, 538, and 54.68—that displayed variability in the variable domains of their light chains and demonstrated a stronger binding to MIN6 cells (Table 1, figure 7, and Figure 1 in supplementary materials). This increased binding was particularly pronounced with Fab_538, as demonstrated by both FACS and IF binding assay analyses. Fab_538 exhibited a stronger binding to MIN6 compared to αTc1 in the IF analysis, with 4 phages binding one MIN6 cell compared to 1,6 binding to one αTc1. This strong binding was also observed with FACS analysis, as 28.8 % of MIN6 cells bound to Fab_538, compared to 8.9 % αTc1 (figures 9 and 10).

Despite the effectiveness of phage display-based screening, unbiased cell-based bio-panning for identifying and characterizing high-affinity, selective, and specific antibodies remains challenging, especially when the expression or prevalence of a cell membrane epitope is downregulated or low (Kim J., 2011). Additionally, certain antigens or epitopes may exhibit low antigenicity or structural variability (Sánchez-Martín D, 2015), which could explain the limited pool of diverse Fabs obtained. In fact, most of the analyzed DNA sequences from the Fab-phages isolated in the fifth round matched the sequence of Fab_53, rather than Fab_538 or Fab_54.68. In the IF binding assays of these three Fabs, we also observed that the binding of Fab_53 to αTc1 cells was stronger than to MIN6 cells, with 2.6 phages binding to a MIN6 cell compared to 4.6 phages binding to an αTc1 cell (figure 2 in the supplementary materials). These findings suggest that the membrane molecule binding Fab_53 is likely expressed by both MIN6 and αTc1 cells, especially in a high-glucose medium.

Another potential reason for the low number of specific and distinct Fabs selected is variability in the amplification of *E. coli* carrying phagemids, along with inconsistent phage display propensities. This could result in limited antigen-antibody binding enrichment across selection rounds, disrupting the enrichment process (Sánchez-Martín D, 2015). As a result, promising binders and newly identified specific Fabs may be lost during successive selection cycles, which could explain why only three distinct Fabs were identified.

Fabs_538, 53, and 54.68 were isolated based on their initial binding to membrane epitopes expressed by MIN6 cells under pathological conditions. To further investigate the impact of glucose concentration on the binding properties of these Fabs, particularly Fab_538, to MIN6 cells, MIN6 cells were cultured for 48 h in 5.5 mM glucose media and then incubated for 2 h with the three Fabs. The binding to MIN6 of all three Fabs increased, with Fab_538 showing the most significant increase in ELISA results (figure 11). These variations suggest that the expression of the MIN6 membrane proteins interacting with the three Fabs, particularly the one binding Fab_538, is influenced by the glucose concentration. The protein P538 seems to be upregulated in media with normal glucose concentrations and downregulated in a high-glucose environment. These findings align with other studies that demonstrated how glucose toxicity contributes to the downregulation of several transcription factors crucial for the expression of proteins involved in maintaining β cell identity and, consequently, their function (Ebrahimi AG, 2020). In fact, it has been shown that even a moderate increase in glucose levels can significantly alter the β cell gene profile in male rats’ pancreas, affecting the expression of several key proteins (Ebrahimi AG, 2020). Many of these proteins are critical for β cell function, including plasma membrane proteins such as the sulfonylurea receptor (Abcc8), the ATP-dependent potassium channel (Kcnj11), and various subunits of the voltage-dependent calcium channel (Cacna1a, Cacna1b, and Cacna1d) (Ebrahimi AG, 2020) (Tuluc P, 2021.) (Flanagan SE, 2009.) (Rodríguez-Rivera NS, 2024.).

Unbiased cell-based Fab-phage display bio-panning provides valuable advantages in identifying novel plasma membrane proteins in their native conformation, along with their corresponding Fabs. However, a key limitation of this method is the time-consuming and expensive process of isolating, purifying, characterizing their structure, and studying the dynamics and functions of the identified Fabs at the experimental level. To address this, we used quicker, relatively reliable, and cost-effective methods to validate and characterize the molecular structure of the selected Fabs, such as AlphaFold, InterProScan, and AF2Complex software. AlphaFold analysis revealed that Fabs_53, 538, and 54.68 had distinct three-dimensional structures, with separate configurations observed between the variable and constant domains of the heavy and light chains (figure 8). Our analysis showed that the three Fabs exhibited variations in specific amino acids within the variable domains of their light chains, distinguishing Fab_53, Fab_538, and Fab_54.68 from one another (Table 1).

InterProScan and AF2Complex tools were subsequently used to predict potential cell membrane epitopes that bind to Fab_538, given its potentially high affinity for MIN6 cells. The InterProScan analysis identified 344 potential *Mus musculus* reactome targets. From these 344 targets, AF2Complex, predicted 650 potential binding proteins for the antigen-binding sites (CDR) on the variable domain of the Fab_538’s light chain. Of these 650 proteins, 10 were selected based on an interface score (IS) greater than 50 %. The proteins analyzed by AF2Complex as potential ligands for Fab_538 were high-quality, experimentally validated proteins corresponding to the 344 epitopes identified by InterProScan. Gao M. et *al.* evaluated the accuracy of the AF2Complex interaction predictions and concluded that IS value of 1 indicates a perfect binding affinity, and values further from 1 suggest a less accurate prediction (Gao M, 2022) Regardless of clear agreement in the literature on specific cutoff intervals for the IS metric generated by AF2Complex and according to Gao M et *al.*’s analysis IS values greater than 0.5 provide fairly reliable prediction ligands.

Among the 10 proteins with an IS greater than 0.5, three are expressed by murine pancreatic β cells, although they have not been identified as part of the cell membrane protein. These include Rab3D, Ap1m2 and Tuba1b. Rab3D, a member of the RAS oncogene family and one of the Rab3 isoforms (including Rab3A, B, and C), plays a crucial role in the secretory process of Golgi-derived vesicles (Fischer von Mollard G, 1994). It is involved in regulating the docking and fusion of these vesicles with the plasma membrane (Fischer von Mollard G, 1994). Several studies have highlighted its role in insulin secretion regulation. For instance, Iezzi et al. demonstrated that the exocytosis of insulin-containing secretory granules can be inhibited by a fraction of all Rab3 isoforms when they are bound to GTP (Iezzi M, 1999).

The role of the Ap1m2 (Adaptor Protein Complex 1 mu 2 subunit) in insulin secretion has not been specifically investigated in pancreatic β cells, but AP-1 implication in insulin secretion and glucose homeostasis in pancreatic β cell was studied and demonstrated *in vivo* by Backes T. M. et *al*. (Backes TM, 2021). Tuba1b (Tubulin, alpha 1b) was also found to be differentially phosphorylated in β cells, suggesting its involvement in insulin secretion within the context of insulin resistance or type 2 diabetes (Mohallem R., 2020).

In order to select internalized Fab-phage clones during rounds, specific strategies are typically used during the selection rounds in order to recover them intact (Mandrup OA, 2017). Since none of our cell-based biopanning rounds involved these strategies, it is possible that intracellular proteins predicted by AF2Complex may not interact with the Fab_538. However, given that Rab3D and Ap1m2 are linked to plasma membrane trafficking, mainly to vesicle docking for Rab3D (Iezzi M, 1999) and receptor-mediated endocytosis for Ap1m2 (Nesterov A, 1999), their spatial localization near the cell membrane increases the likelihood that they could be potential binding proteins for Fab_538.

Among the seven other predicted binding proteins for Fab_538, Trac, Opn1sw, Cd8a, and Tas2r41 have been characterized as transmembrane proteins expressed in cell lines other than pancreatic β cells in previous studies. Trac (T Cell Receptor Alpha Constant) is primarily expressed by T lymphocytes (Peters LD, 2023). While the interaction between pancreatic β cells and T lymphocytes has been studied in the context of type 1 diabetes, no studies have shown that β cells themselves express Trac (Peters LD, 2023). Opn1sw (Opsin 1, Short Wave Sensitive) is a plasma membrane protein expressed in retinal cone photoreceptor cells. Similarly to Trac protein, no studies have confirmed its expression in pancreatic β cells. Nonetheless, optogenetic approaches have been employed to investigate the hormonal secretions and functions of pancreatic islets (Gangemi CG, 2024).

CD8a (CD8 subunit alpha), a membrane protein, is an important component of CD8+ T cell receptor complex and the phenotypic marker for cytotoxic T cells (Raskov H, 2021). Cytotoxic T cells are implicated in several cancers and autoimmune diseases, including type 1 diabetes (Gearty SV, 2022). In T1D, insulin-producing cells are targeted and eliminated by the β cell-specific CD8+ T cells, but the triggering mechanisms of these events are still under investigation (Gearty SV, 2022). Tas2r41 (taste receptors member 2, 41) is also a plasma membrane protein, although its expression in insulin-secreting β cells has not been established. However, the expression of the receptors TAS2R has been described in the human tongue, precisely at the taste buds, and they are associated with bitter taste sensations (Lipchock SV, 2013.). The potential link between insulin secretion and bitter taste receptors was investigated in one study that showed that the activation of TAS2R38 receptors can stimulate the secretion of glucagon-like peptide-1 (GLP-1) by enteroendocrine L-cells (Pham H., 2016). Interestingly enough, because TAS2R38 expression has been observed in CD8+ T cells, its role in immune modulation responses has been investigated, however not specifically in T1D (Tuzim K., 2021).

AF2Complex predictions for ligands provided us with a very interesting list of potential ligand candidates, however it would be valuable to explore the interaction between Fab_538 and these predicted ligand proteins in future experiments.

## Conclusion

Monoclonal antibody production and the experimental processes for identifying new disease-modifying drugs are highly demanding, which calls for more economical approaches to address these challenges. This study serves as a proof of concept, showing that unbiased screening for new cell membrane proteins can be made more time-efficient and cost-effective by utilizing cell-based biopanning with Fab-phage display technology, combined with bioinformatics tools like InterProScan, AlphaFold and AF2Complex.

