## supplements/645157_file05.pdf for "Investigating pancreatic β cell membrane epitopes using unbiased cell-based Fab-phage display"

### Suplementary materials

##### Protein alignment of the VL and CL regions between Fabs_538 and 53

**a.**

```
Matrix: EBLOSUM62
Gap penalty: 2.0
Extend penalty: 2.0
Score: 1075.0
Fab_53_VL+CL length:213
Fab_538_VL+CL length:216
Alignment length: 216
Identity: 207/216 (95.83%)
Similarity: 211/216 (97.69%)
Gaps: 3/216 (1.39%)

Fab_53: 1   MTQSPISLPVTPGEPASISCRSSQSLLFSNGYNYLDWYLQKPGQSPQLLIYLGSNRASGV 60
Fab_538:1   MTQSPLSLPVTPGEPASISCRSSQSLLHSNGYNYLDWYLQKPGQSPQLLIYLGSNRASGV 60

Fab_53: 61  PDRFSGSGSGTDFTLQISRVEAEDVGVYYCMQALQTPLTFGGGTKVEIKRTVAAPSVFIF120
Fab_538:61  PDRFSGSGSGTDFTLKISRVEAEDVGVYYCMQALQTPLTFGQGTRLEIKRTVAAPSVFIF120

Fab_53: 121 PPSDEQLKSGTASVVCLLNNFYPREAKVQWKVDNALQSGNSQESVTEQDSKDSTYSLSST180
Fab_538:121 PPSDEQLKSGTASVVCLLNNFYPREAKVQWKVDNALQSGNSQESVTEQDSKDSTYSLSST180

Fab_53: 181 LTLSKADYEKHKVYACEVTHQGLSSPVTKSFNR---213
Fab_538:181 LTLSKADYEKHKVYACEVTHQGLSSPVTKSFNRGEC216
```

#### Protein alignment of the VL and CL regions between Fabs_538 and 54.68

**b.**

Gap penalty: 2.0
Extend penalty: 2.0
Score: 1085.0
Fab_538_VL+CL length:216
Fab_54.68_VL+CL length:213
Alignment length: 216 Identity: 209/216 (96.76%)
Similarity: 211/216 (97.69%)
Gaps: 3/216 (1.39%)

```
Fab_538:     1 MTQSPLSLPVTPGEPASISCRSSQSLLHSNGYNYLDWYLQKPGQSPQLLIYLGSNRASGV 60
Fab_54.68:   1 MTQSPLSLPVTPGEPASISCRSSQSLLHSNGYNYLDWYLQKPGQSPQLLIYLGSNRASGV 60

Fab_538:    61 PDRFSGSGSGTDFTLKISRVEAEDVGVYYCMQALQTPLTFGQGTRLEIKRTVAAPSVFIF120
Fab_54.68:  61 PDRFSGSGSGTDFTLKISRVEAEDVGVYYCMQALQAPLTFGGGTKVEIKRTVAAPSVFIF120

Fab_538:   121 PPSDEQLKSGTASVVCLLNNFYPREAKVQWKVDNALQSGNSQESVTEQDSKDSTYSLSST180
Fab_54.68:121 PPSDEQLKSGTASVVCLLNNFYPREAKVQWKVDNALQSGNSQESVTEQDSKDSTYSLSST180

Fab_538:   181 LTLSKADYEKHKVYACEVTHQGLSSPVTKSFNRGEC216
Fab_54.68:181 LTLSKADYEKHKVYACEVTHQGLSSPVTKSFNR---213
```

#### Protein alignment of the VL and CL regions between Fabs_53 and 54.68

**c.**

Gap penalty: 2.0
Extend penalty: 2.0
Score: 1084.0
Fab_53_VL+CL length:213
Fab_54.68_VL+CL length:213
Alignment length: 213
Identity: 209/213 (98.12%)
Similarity: 211/213 (99.06%)
Gaps: 0/213 (0.00%)

```
Fab_53:      1 MTQSPISLPVTPGEPASISCRSSQSLLFSNGYNYLDWYLQKPGQSPQLLIYLGSNRASGV60
Fab_54.68:   1 MTQSPLSLPVTPGEPASISCRSSQSLLHSNGYNYLDWYLQKPGQSPQLLIYLGSNRASGV60

Fab_53:     61 PDRFSGSGSGTDFTLQISRVEAEDVGVYYCMQALQTPLTFGGGTKVEIKRTVAAPSVFIF120
Fab_54.68:  61 PDRFSGSGSGTDFTLKISRVEAEDVGVYYCMQALQAPLTFGGGTKVEIKRTVAAPSVFIF120

Fab_53:    121 PPSDEQLKSGTASVVCLLNNFYPREAKVQWKVDNALQSGNSQESVTEQDSKDSTYSLSST181
Fab_54.68: 121 PPSDEQLKSGTASVVCLLNNFYPREAKVQWKVDNALQSGNSQESVTEQDSKDSTYSLSST181

Fab_53:    180 LTLSKADYEKHKVYACEVTHQGLSSPVTKSFNR213
Fab_54.68: 180 LTLSKADYEKHKVYACEVTHQGLSSPVTKSFNR213
```

**Figure 1: a.)** Comparative amino acid sequence analysis of the VL+CL domains between Fab_538 and Fab_53**. b.)** Comparative amino acid sequence analysis of the VL+CL domains between Fab_538 and Fab_54.68**. c.)** Comparative amino acid sequence analysis of the VL+CL domains between Fab_53 and Fab_54.68**. These analyses were performed using Vector Builder website: https://en.vectorbuilder.com/.**

##### Immunofluorescence results of the Fab-phage_53

a.

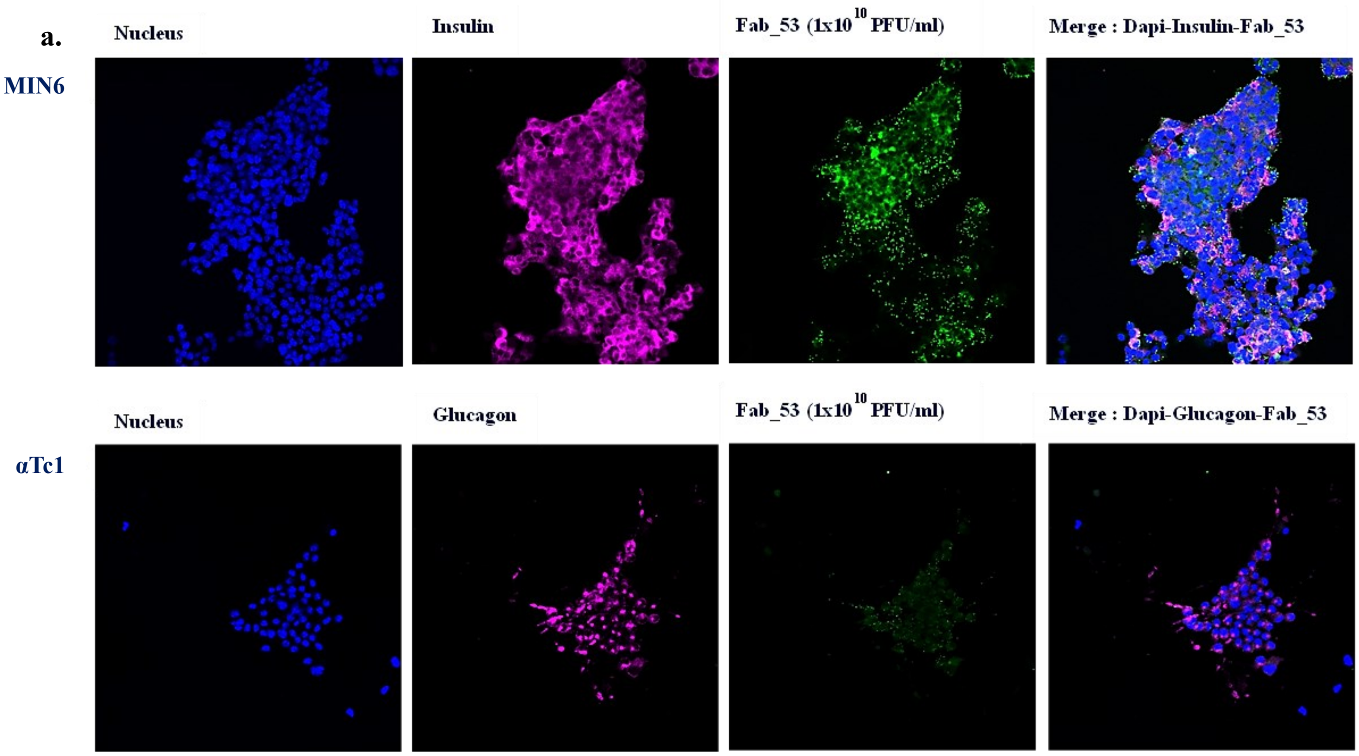


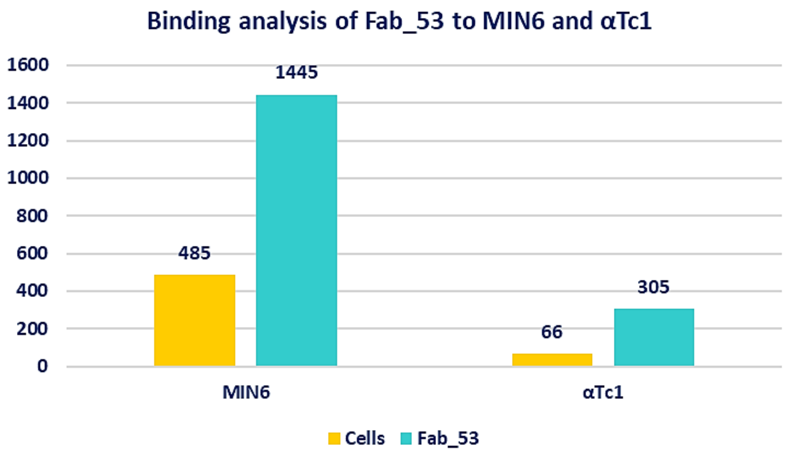


**Figure 2: a.)** The binding of Fab-phage_53 to MIN6 cells was confirmed through immunofluorescence. Blue represents DAPI staining for the nucleus, purple indicates Alexa 647 staining for insulin in MIN6, glucagon in αTC1. Green represents Alexa 488 staining for Fab_53 in MIN6 and αTC1 cells. **b.)** Fab-phage_53 binding unit to individual cell types and cell count per image were estimated by using ImageJ intensity analysis software. It was confirmed that Fab-phage_53 can bind to both MIN6 and αTc1, with 1445 phages binding to 485 cells and 305 phages binding to 66 αTc1 cells. The binding rate for αTc1 is 4.6 times and for MIN6 it is around 3.

#### Analysis of the variable and constant domains of the light and heavy chains of Fabs, 538, 53, and 54.68 using interProScan

##### Fab_538

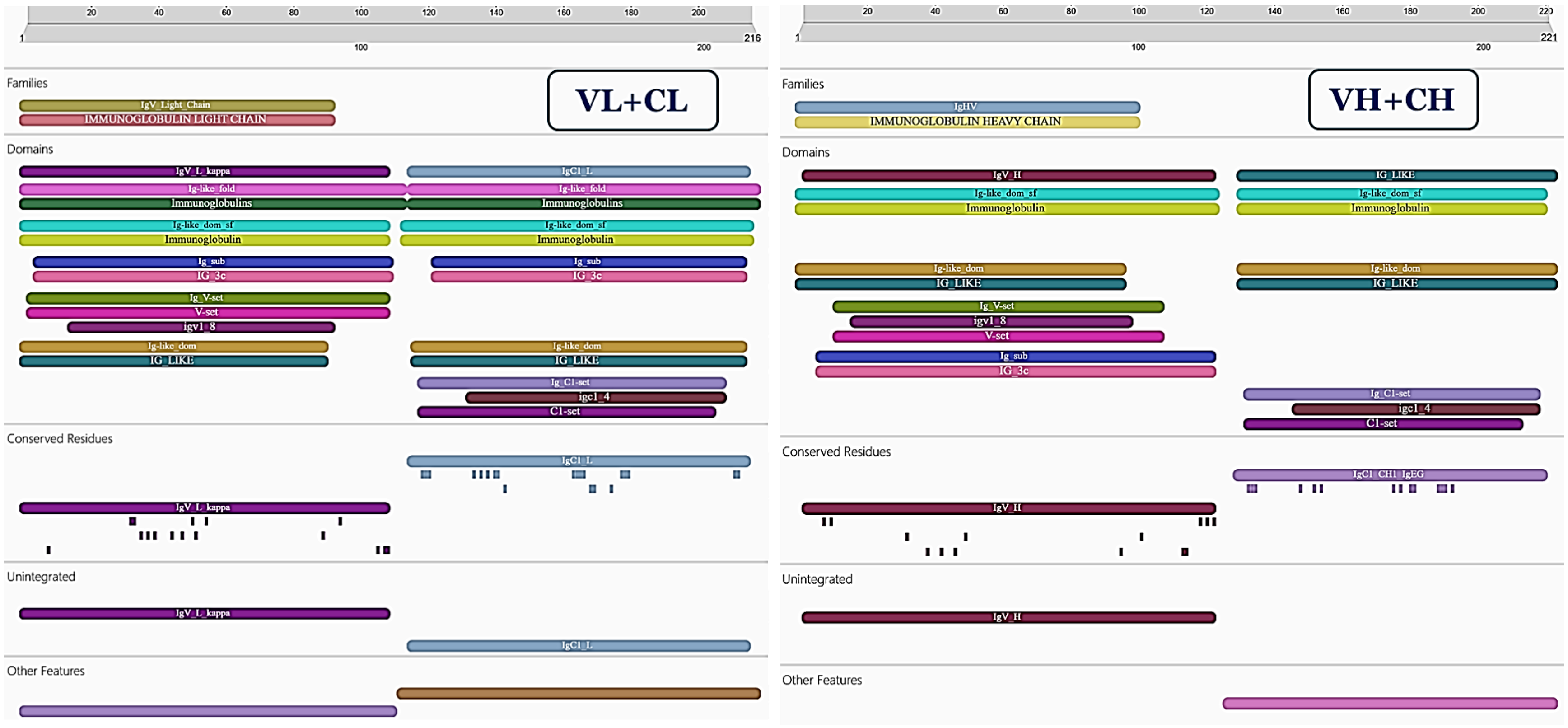


**Figure 3:** The amino acid sequences of Frame 1 for the VL, CL, VH, and CH of Fab_538 correspond to the sequences of an IgG class antibody, with the light chain belonging to the kappa group. https://www.ebi.ac.uk/interpro/search/sequence/

#### Fab_53

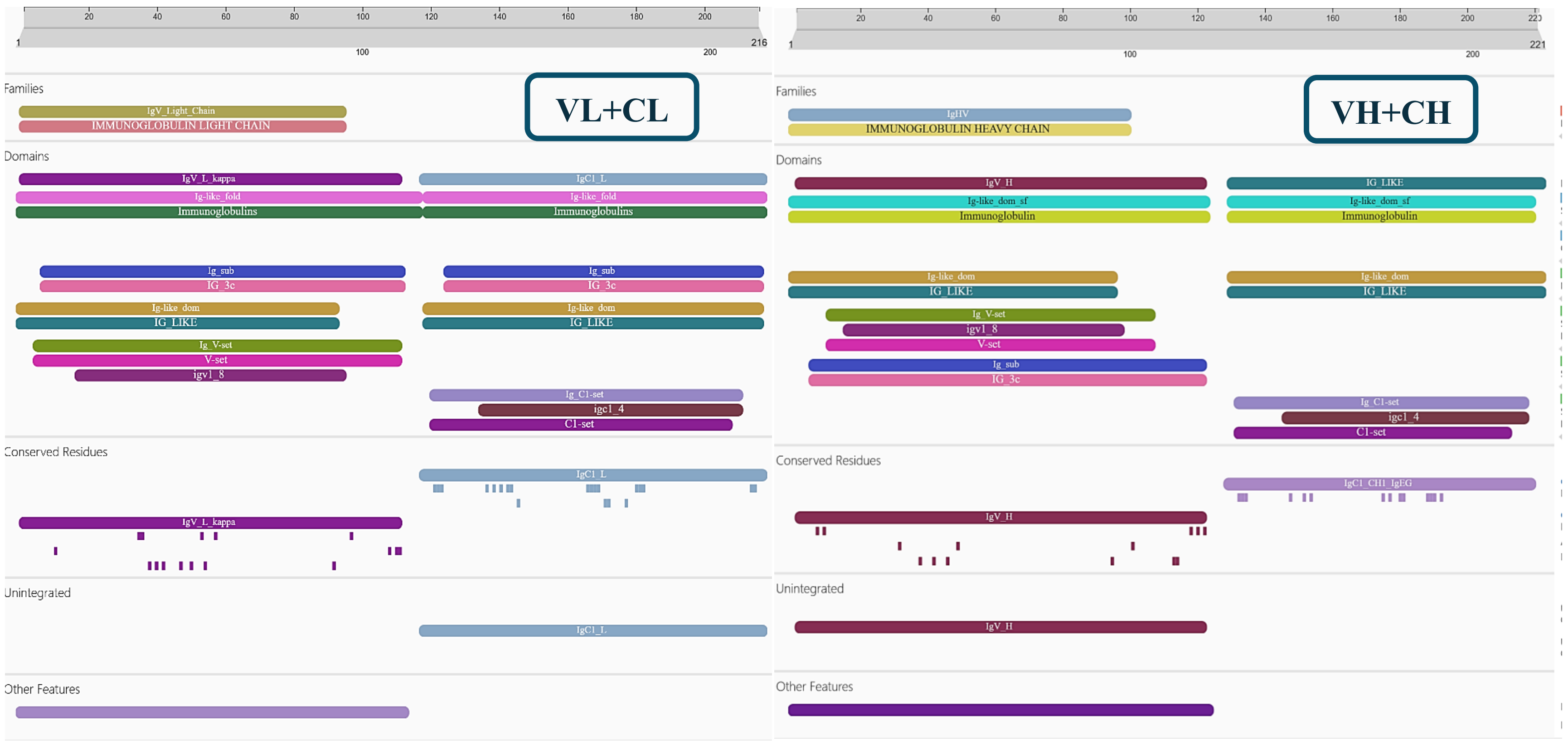


**Figure 4:** The amino acid sequences of Frame 1 for the VL, CL, VH, and CH of Fab_53 correspond to the sequences of an IgG class antibody, with the light chain belonging to the kappa group. https://www.ebi.ac.uk/interpro/search/sequence/

#### Fab_54.68

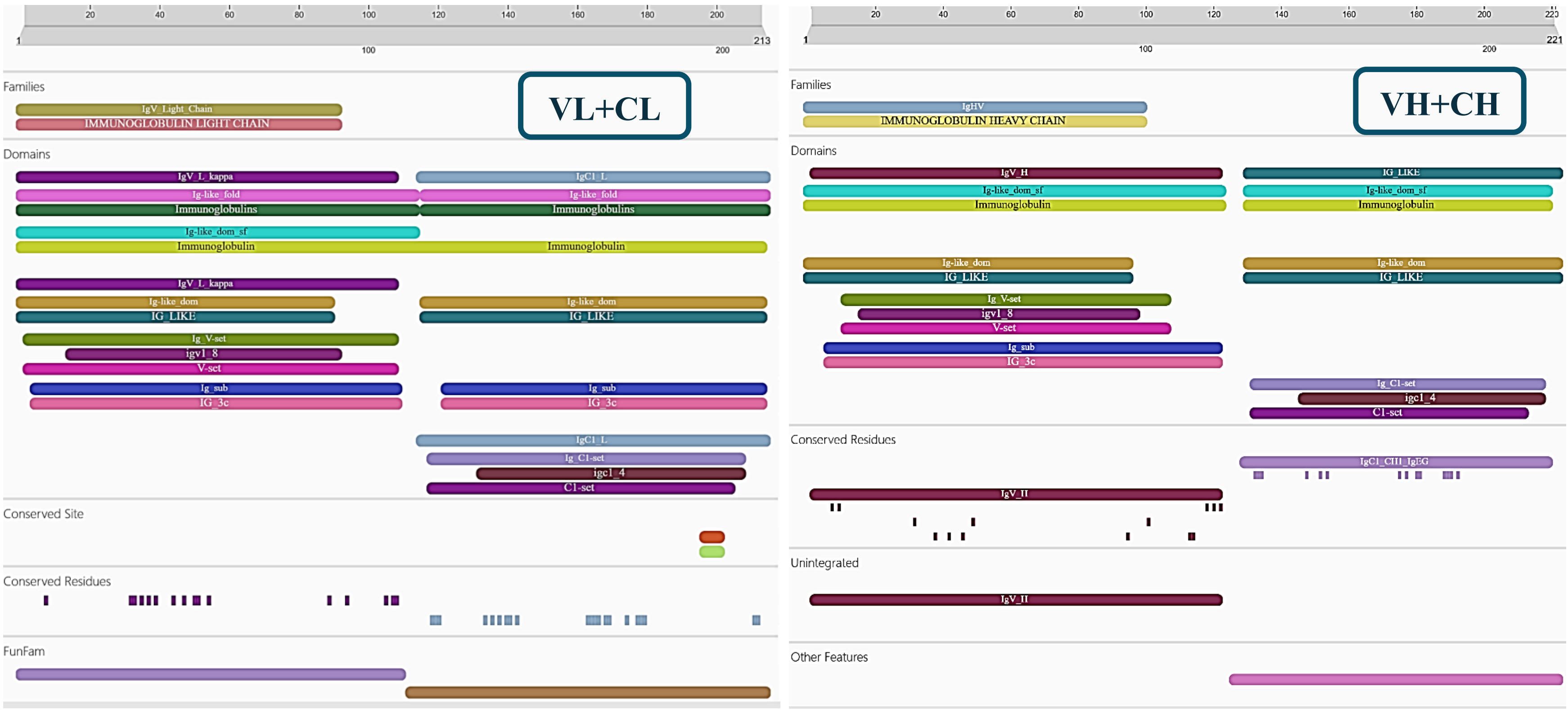


**Figure 5:** The amino acid sequences of Frame 1 for the VL, CL, VH, and CH of Fab_54.68 correspond to the sequences of an IgG class antibody, with the light chain belonging to the kappa group. https://www.ebi.ac.uk/interpro/search/sequence/

| Fab_538 | Light chains lenght | |
|---|---|---|
| | V | C |
| Ig Variable Light Chain | 1-92 | |
| Ig C1-set | | 117-203 |
| Ig-like domain | 1-90 | 115-212 |
| Ig-like fold | 1-113 | 114-213 |
| Ig V-set domain | 15-92 | |
| Ig domain subtype | 5-109 | 121-212 |
| Ig-like domain superfamily | 1-108 | 112-214 |
| Ig/major histocompatibility complex, conserved site | | 194-200 |

**Table 1: InterProScan analysis results of Fab_538.** InterProScan analysis of Fab_538 reveals that the amino acids in Frame 1 of the VL+CL antigen-binding sites are identical to those found in IgG.

| | Temperatures in Celsius | Time |
|---|---|---|
| Hot start-1 | 94 | 240 seconds (4 min) |
| Denaturation-2 | 94 | 30 sec |
| Annealing-3 | 61 | 30 sec |
| Elongation-4 | 72 | 150 sec (2,30 min) |
| Repetion of 2 to 4 | 29x | |
| Final elongation | 72 | 900 sec (15 min) |
| Hold | 4 | ∞ |

**Table 2:** PCR cycling parameters programme of Fab-DNA amplification.

**Table 3**: List of commercial antibodies used in FACS, ELISA and IF

| Antibody | Type | Company | Cat # | Dilution |
|---|---|---|---|---|
| Anti-M13 Major Coat Protein (RL-ph1) | Mouse mAb | Santa cruz biothechnology, INC. | Sc-53004 | 1:1000 |
| Anti-Insulin | Rabbit mAb | Cell Signaling Technology | C27C9 | 1:250 |
| Anti-Glucagon | Rabbit pAb | Cell Signaling Technology | 2760 | 1:500 |
| Anti-Transferrin Receptor (Cd71) | Rabbit pAb | abcam | Ab84036 | 1ug/ml |
| Anti-Cd99 | Rabbit pAb | Invitrogen-ThermoFisher scientific | PA5-102556 | 1:200 |
| Wheat Germ Agglutinin (WGA) | Conjugated to Alexa Fluor 488 | Invitrogen- Thermo Fisher Scientific | W11261 | 5ug/ml |
| Alexa 488 | Donkey anti mouse IgG | Invitrogen- Thermo Fisher Scientific | A-21202 | 1:500 |
| Alexa 647 | Donkey anti-rabbit IgG | Invitrogen- Thermo Fisher Scientific | A31573 | 1:500 |
| Alexa 647 | Goat anti-mouse IgG | Invitrogen- Thermo Fisher Scientific | A21235 | 1:500 |
